# Reproductive compensation and selection among viable embryos drive the evolution of polyembryony

**DOI:** 10.1101/2020.11.17.387340

**Authors:** Yaniv J Brandvain, Alexander J Harkness, Tanja Pyhäjärvi

## Abstract

Simple polyembryony – where one gametophyte produces multiple embryos with different sires but the same maternal haplotype – is common among vascular plants. We show that together polyembryony’s two benefits – “reproductive compensation” achieved by providing a backup for inviable embryos, and the opportunity to favor the fitter of surviving embryos, can favor the evolution of polyembryony. To do so, we develop an infinite-site, forward population genetics model to test how these factors can favor the evolution of polyembryony, and how these underlying benefits of polyembryony shape the genetic load under a range of biological parameters. While these two benefits are difficult to disentangle in nature, we construct variant models of polyembryony that either only include or only exclude the opportunity for reproductive compensation. We find that reproductive compensation strongly favors the evolution of polyembryony, and that polyembryony is favored much more weekly in its absence, suggesting that the benefit of a backup embryo is the force favoring polyembryony. Remarkably we find nearly identical results in cases in which mutations impact either embryo or post-embryonic fitness (no pleiotropy), and in cases in which mutations have identical fitness effects embryo or post-embryonic fitness (extreme pleiotropy). Finally, we find that the consequences of polyembryony depends on its function – polyembryony results in a decrease in mean embryonic fitness when acting as a mechanism of embryo compensation, and ultimately increases mean embryonic fitness when we exclude this potential benefit.

Nature is, above all, profligate. Don’t believe them when they tell you how economical and thrifty nature is.

– Annie Dillard 1974.

---

Not only do most parents produce more offspring than will survive, but most organisms that provide parental care make more offspring than they will likely be able to nurture to independence. Frequent siblicide in the great egret, *Casmerodius albus*, provides a dramatic example of this – siblings kill one another, presumably over the ability to monopolize small food items (Mock 1984); why then do egret mothers continue laying eggs that will develop into offspring that will kill one another? Could such overproduction allow parents to preferentially raise high fitness offspring for offspring quality (Forbes and Mock 1998), or does the “diverse portfolio” of offspring born over the breeding season allow parents to hedge their bets, ensuring survival of at least one chick (Forbes 2009)?

Simple polyembryony – in which a single maternal gametophyte is fertilized by multiple sperm cells to produce multiple embryos with genetically identical maternally derived genomes but distinct paternal genomes (Buchholz 1922; Schnarf 1937, cited in Dogra 1967) – provides an even more extreme, but perhaps less dramatic, example of this problem. Simple polyembryony is ubiquitous in gymnosperms (Willson and Burley 1983), and is found in many seedless vascular plants including ferns and horsetails (Buchholz 1922). The number of archegonia per seed typically varies from two to four in the genus *Pinus*, but can reach up to 200 (as reported in *Widdringtonia juniperoides* Saxton 1934), of which only a single single embryo typically survives in mature seed (Chamberlain 1966). We develop a population genomic model to both test the two major advantages of simple polyembryony described by Kärkkäinen and Savolainen (1993), *reproductive compensation* (akin to egret mothers hedging their bets), and *fitter offspring* (akin to egret mothers preferentially raising high-fitness offspring) theories for the evolution of polyembryony, and to investigate how polyembryony changes embryonic and post-embryonic fitness and the genetic architecture of these complex traits. While these benefits of polyembryony have been acknowledged for some time, they have rarely been discussed together and their theoretical contributions to the evolution of polyembryony have not been disentangled (but see Ferreira et al. 2019, for an empirical study of the influences of these processes on seedling establishment in a population of *Carapa surinamensis* that produces mono- and polyembryonic seeds).

Reproductive compensation is an increase in seed set that occurs when embryo mortality is counteracted by an expanded supply of embryos (Sorensen 1982; Porcher and Lande 2005). Polyembryony provides reproductive compensation if a lone embryo is less likely to develop into a successful seed than is a collection of sibling embryos. So, for example, if a proportion *p* of embryos are inviable, and the survival of each embryo is independent, a second embryo increases the probability that a seed contains a surviving embryo from 1 *− p* to 1 *− p*^2^ (Lindgren 1975).

Alternatively, polyembryony could be favored because it allows for the preferential development of the fitter offspring. This form of ” Developmental Selection” (see Buchholz (1922)) would be particularly beneficial if embryonic and post-embryonic fitness are positively correlated. Such a correlation can arise either through pleiotropy (embryonic and post-embryonic fitness effects of the same mutation) across the life cycle, or if embryonic fitness determined by one set of loci predicted post-embryonic fitness produced by another set of loci. This later option seems particularly likely if inbred offspring are unfit across the life cycle, and as such, simple polyembryony is often interpreted as an inbreeding avoidance mechanism (e.g. Dogra 1967; Sorensen 1982) analogous to the self-incompatibility systems (hereafter SI) found in angiosperms. Koski (1971) and others contend that this gives way to evolution of the so-called “Embryo Lethal System” – an apparently coordinated self destruction mechanism revealed upon inbreeding (or the production of low-vigor embryos, more generally) in pines as a mechanism evolved to prevent selfing (Koski 1971; Sarvas 1962, e.g. page 162 onwards). Under this model, polyembryony does not prevent self-fertilization *per se*, but dampens self-fertilization’s deleterious effects by allowing selection between embryos, which would favor outcrossed over inbred progeny before major maternal resource allocation (Willson and Burley 1983; Sorensen 1982). As such this model could circumvent the constraint imposed by the unenclosed gymnosperm seed, which precludes prezygotic mate choice (e.g. SI systems Dogra 1967; Sorensen 1982; Willson and Burley 1983).

We note that there is an inherent difficulty in distinguishing between the reproductive compensation model (’picking a surviving offspring’) and the fitter offspring model – as clearly a surviving offspring has a higher fitness than and would be preferable to an inviable offspring. As such, reproductive compensation is somewhat baked into the fitter offspring model. To disentangle these benefits of polyembryony we consider alternative models for its evolution. First we introduce a model with both such benefits of polyembryony – that is, if both embryos survive viability selection, the embryo that makes it to the seed is chosen in proportion to its relative fitness. We then model the benefit of reproductive compensation by randomly choosing among the embryos that survive viability selection – thus if one embryo does not survive viability selection, the mother is left with a backup. Finally we introduce a control that separates the benefits of having a higher fitness offspring from the benefit of reproductive compensation by allowing for selection between embryos only when both survive, and choosing an embryo randomly otherwise. Our ability to distinguish the benefit of reproductive compensation the benefit of ”selection between surviving embryos” (as we label the model hereafter) comes at a cost of biological realism, and we warn the reader that this control variant of our model is presented to guide our thinking, not to model biological reality.

Critically, fitter offspring model assumes that possibility of effective mechanism by which the fitness of embryos in a seed can be differentiated, a topic of much debate. Based on extensive experimental work on *P. sylvestris*, Sarvas (1962) stated that the ”struggle for life” is quite apparent under microscopic observation of developing embryos, but unfortunately the details leading to this conclusion were not disclosed. Empirical studies evaluating these ideas are quite rare, and the evidence from these studies is mixed. Most such studies address the possibility of selection between surviving embryos by comparing the observed frequency of selfed embryos (which are likely to be low-fitness) to their expectation under a model in which embryos are chosen at random. In a particularly careful study, O’Connell and Ritland (2005) conducted controlled pollination with varying levels of self-pollen with *Thuja plicata*, and found that the effect of embryo competition became apparent with a probability of selfing (0.75), that exceeds reasonable estimates of the frequency of self-pollination in most conifers. While subtle effects at lower selfing rates are plausible, other researchers argue that selfed embryo death primarily occurs after the dominant embryo is determined (Williams 2008), and embryo survival is determined by chance physical factors (Williams 2007; Mikkola 1969), undercutting the fitter offspring model.

In addition to various selective forces favoring the evolution of polyembryony, polyembryony itself can have striking evolutionary consequences. Previous models have investigated how polyembryony (or less mechanistically explicit forms of reproductive compensation) could shape the genetic load (Latta 1995; Sakai 2019; Porcher and Lande 2005; Kärkkäinen et al. 1996) and the exposure of inbreeding depression (Kärkkäinen and Savolainen 1993; Hedrick et al. 1999). These models generally show that by removing selfed embryos early in development, polyembryony will prevent the effective purging of deleterious recessive mutations (Klekowski 1982; Haig 1992), and will therefore increase the number of deleterious mutations at equilibrium, and increase the extent of inbreeding depression. Despite its weakening effect on selection, in such models polyembryony also decreases the realized selfing rate, and ultimately increase population mean fitness. As such, polyembryony is often suggested as an explanation for the joint observation of high inbreeding depression (gymnosperms have an estimated 5-10 lethal equivalents per haploid genome (Lynch and Walsh 1998; Williams and Savolainen 1996)), the low realized selfing rates in gymnosperms (Kärkkäinen and Savolainen 1993; Hedrick et al. 1999), and the absence of a relationship between inbreeding depression and the primary selfing rate in gymnosperms (Husband and Schemske 1996). However other models of polyembryony make drastically different predictions — for example, Latta (1995) modelled the fitter embryo component of polyembryony (which he called ”embryo competition”) and found that polyembryony increased the efficacy of selection and decreased the number of deleterious mutations per individual. Here, we find that much of these differences are attributable to implicit, and often unacknowledged, modelling decisions that consider polyembryony as a mechanism of embryo compensation or selection between surviving embryos – that is, when polyembryony functions as a mechanism of embryo compensation we find an elevated embryo load because selection is relaxed (as found in e.g. Lande et al. 1994; Klekowski 1982; Haig 1992), while when functioning as a mechanism of selection between viable embryos, polyembryony tends to more effectively remove the mutational load of embryos (as found in Latta 1995).

Previous research provides some insight into the evolutionary consequences of polyembryony, but contains numerous modelling assumptions that limit their applicability to major questions in the evolution of polyembryony and its consequences. For example, comparing cases with and without reproductive compensation, Porcher and Lande (2005) showed that reproductive compensation can favor the evolution of selfing and can allow for the maintenance of mixed mating systems, while Sakai (2019) showed that selective embryo abortion could allow for the maintenance of high levels of inbreeding depression in selfing species. But if the mating system of an initially monoembryonic population affects whether polyembryony evolves in the first place, this initial condition may affect subsequent mating system evolution after the transition to polyembryony. A second limitation with current theory of the evolutionary consequences of polyembryony is that each model has focused on a single dominance and selection coefficient. As such, while current theory predicts evolution of the number and frequency of deleterious mutations, the magnitude of genetic load, it cannot predict evolution of the distributions of dominance or selection coefficients, or the architecture of genetic load. This limitation has prevented theory from addressing Koski’s (1971) hypothesis that the “Embryo Lethal System” evolved as an altruistic mechanism by which inbred embryos sacrifice themselves to prevent their mothers from selfing, as opposed to the parsimonious alternative that selfing simply exposes the elevated number of deleterious mutations that can accumulate under polyembryony.

Here we present a series of infinite-sites forward population genetic simulations of polyembryony to evaluate how alternative benefits of polyembryony favor its evolution and shape the genetic load. We limit our focus to “simple polyembryony” and do not consider cleavage polyembryony, in which a fertilized zygote can split into numerous genetically identical embryos (Agapito-Tenfen et al. 2012), or nucellar polyembryony, in which maternal tissue asexually develops into embryos (Lakshmanan and Ambegaokar 1984), sometimes competing with sexually derived embryos, as they are likely favored by distinct mechanisms (Ganeshaiah et al. 1991).

## Methods

### Overview

We present a series of models to disentangle the contribution of the potential evolutionary benefits of reproductive compensation and favoring the fitter embryo to the evolution of polyembryony. To better understand how and when these factors favor the evolution of polyembryony, we vary the distribution of dominance and fitness effects and the probability of selfing. Finally, when polyembryony does evolve, we ask how its evolution shapes the genetic load and its architecture.

The life cycle begins with *N* = 1000 diploid seeds, each of which has one or two embryos, depending on whether mothers are mono- or polyembryonic. Following embryo selection (which can include viability selection and/or selection between embryos – see below), surviving seed parents for the next generation are chosen with replacement with a probability reflecting their post-embryonic fitness. Each time a seed parent is chosen, it generates one of the *N* seeds in the next generation, thus maintaining a constant population size of *N* viable plus inviable seeds every generation. Each embryo in the seed is fertilized independently. If directly selfed, which occurs with probability equal to the selfing rate, the seed parent of an embryo is also its pollen parent, otherwise the pollen parent is selected at random and in proportion to adult fitness. Next, gametes are formed by free recombination and each gamete acquires mutations. Finally, fusion between gametes generates the embryos for the next generation (Fig. 1).

**Figure 1:**
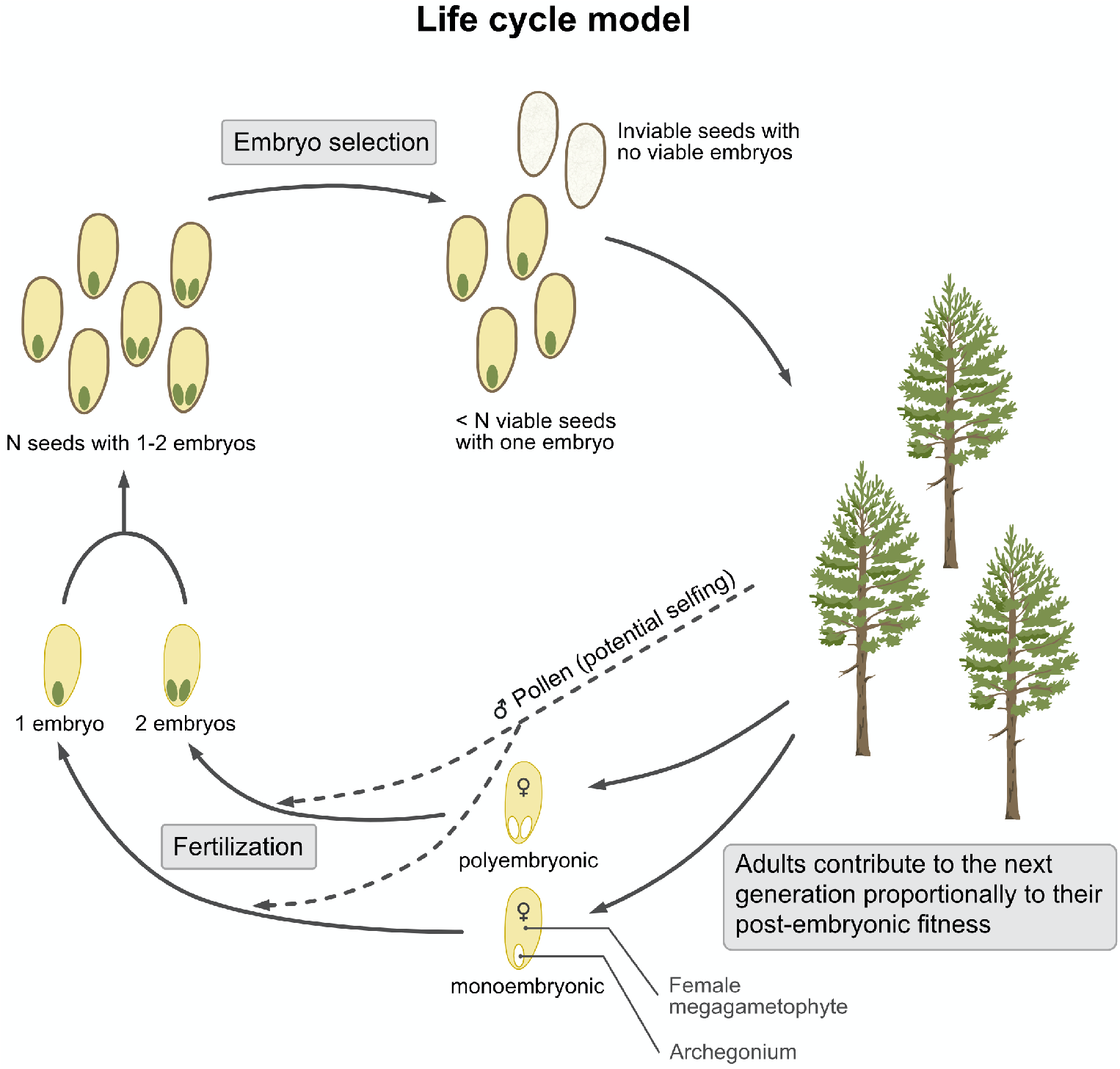
Overview of the life cycle model: The life cycle starts with *N* seeds, each with one or two embryos, followed by embryo and seed selection. Because of seed inviability, the number of plants will be less than *N*. Seed parents are chosen with replacement in proportion to their post-embryonic fitness, and directly self with probability *p*_self_. Both embryos of polyembryonic mothers are fertilized independently, and pollen parents of non-selfed seed are sampled with from the population with replacement in proportion to each genotypes post-embryonic fitness. Seed parents carrying the dominant polyembryony allele, produce two archegonia per seed, while those without this allele produce only one.

### Parameters and model details

#### Genome structure and mutation rate

Every generation, each haploid genome expects a Poisson distributed number (mean *U*) of *de novo* deleterious mutations at any one of an infinite number of unlinked sites (i.e. an infinite sites model). We investigate cases with *U* = 0.5 mutations per haploid chromosome per generation. This value is in the middle of estimates of the per generation haploid mutation rates in small-statured plants (with ranges of 0.05 to 2.0 as reviewed by Schultz and Scofield (2009) where *U* is presented as the diploid rate, such that values are twice ours). While this value could differ in large-statured long-lived plants (Schultz and Scofield 2009), initial explorations of numerous values of *U* led to qualitatively similar results (so long as populations didn’t crash under their mutational load).

#### The timing of mutational effects

We focus on the case in which half of *de novo* deleterious mutations impact embryonic fitness and the other half impact post-embryonic fitness. While our main results ignore the possibility that mutations may impact embryonic and post-embryonic fitness, our code also allows mutations to influence both embryonic and post-embryonic fitness pleiotropically, and we provide results for a special case of ”complete pleiotropy” in the discussion.

#### The distribution of fitness and dominance effects of new mutations

For all parameter values, fitness effects, *s*, across loci are multiplicative and independent such that the fitness of the *i*^*th*^ individual, *w*_*i*_, equals the product of one minus the deleterious effect of their genotype at the *k*^*th*^ locus, taken across all loci (i.e. *w*_*i*_ = Π_*k*_(1 *− s*_*ik*_)). To investigate the impact of mutational architecture on the evolution of polyembryony, we compare models with a different value of fitness and dominance effects of new mutations. For s we present cases with *s* = 0.1, *s* = 0.5, *s* = 1, and *s* = Uniform(20*/N*, 1). Dominance, *h*, can take any value between 0 and 1, but we present cases with full recessivity (*h* = 0) and full additivity (*h* = 0.5), as well as a case where the dominance of each mutation takes a random value between zero and one from the uniform distribution (*h* = Uniform(0, 1)). Thus, mutational effects span the range from quite deleterious to lethal, but will not reach fixation by random genetic drift. Practically, this means that we save considerable computational resources, and that we do not consider weakly deleterious mutations whose fixation is not effectively prevented by selection. In all simulations we assumed that the distribution of fitness and dominance effects did not differ for mutations impacting the embryo and adult.

#### Selfing

With a probability equal to *p*_self_ (which we systematically varied from zero to one in increments of 0.2) the seed parent was also chosen to be the pollen parent. Otherwise, pollen parents chosen at random and with replacement in proportion to adult fitness, using the sample() function in R (R Core Team 2020). We note that this random mating does not preclude selfing. Therefore, even with *p*_self_ = 0, one of every *N*_*e*_ embryos (approximately 0.001 when N = 1000, depending on seed survival rates and the variance in post-embryo fitness) is expected to have identical pollen and seed parents.

### Evolution

#### Burn-in

For all parameter combinations, we forward-simulated ten replicate processes for 2000 generations, ensuring that populations achieved mutation-selection-drift balance by visually examining the variability in the number of deleterious mutations over time and among replicates (Figure S1). For most parameter values, equilibrium was reached within this time frame (Figure S1). However, for recessive mutations in predominantly outcrossing populations (with selfing rates of 0, 0.2, or 0.4) this was not enough time to reach equilibrium. For these slowly equilibrating cases, we increased the burn-in period until 3000 generations, at which point equilibrium was largely achieved. However, with complete selfing, a non-recessive load, and *s* = 0.1, the number of deleterious mutations seems somewhat unstable (that is, under these parameter values, deleterious mutations with *s* = 0.1 eventually reach fixation, and therefore the number of mutations per individual eventually increases – Figure S1). To ensure that our simulations performed sensibly, we compared results from these burn-ins into analytical expectations when known, specifically focusing on the expected number of recessive alleles per individual (Gao et al. 2015).

#### Invasion of polyembryony

For each burn-in replicate, we ran many introductions of a dominant acting polyembryony allele, introduced at a frequency of 1/2*N*, and kept track of the fate of this allele (loss or fixation) for each introduction. Due to computational considerations, we varied the number of introductions from 500 to 1000 for each model of polyembryony (below) for each burn-in replicate. That is, when polyembryony was strongly favored, a given simulation took longer to complete (because fixation from 1/2*N* takes more time than loss from 1/2*N*). By contrast, when polyembryony is not strongly favored, individual simulations are faster (because loss occurs more quickly than fixation) and more precision was needed to distinguish fixation rates from neutrality. The R (R Core Team 2020) code for these forward simulations is available as a supplementary document.

### Models of polyembryony

We aim to dissect the contribution of reproductive compensation and the benefit of raising a higher fitness seed to the evolution of polyembryony.

A benefit of having a higher fitness embryo induced by polyembryony is that, mothers can increase their expected inclusive fitness by allowing for selection between embryos if doing so results in higher fitness seeds (Bottom panel of Figure 2A). We allow for the benefit of having a higher fitness seed by selecting the embryo that makes it to the seed among surviving embryos with probabilities in proportion to their embryonic fitness. We can effectively remove this benefit of polyembryony by randomly choosing one among viable embryos in a seed to become the dominant embryo and continue development. This model isolates the benefits of reproductive compensation.

**Figure 2:**
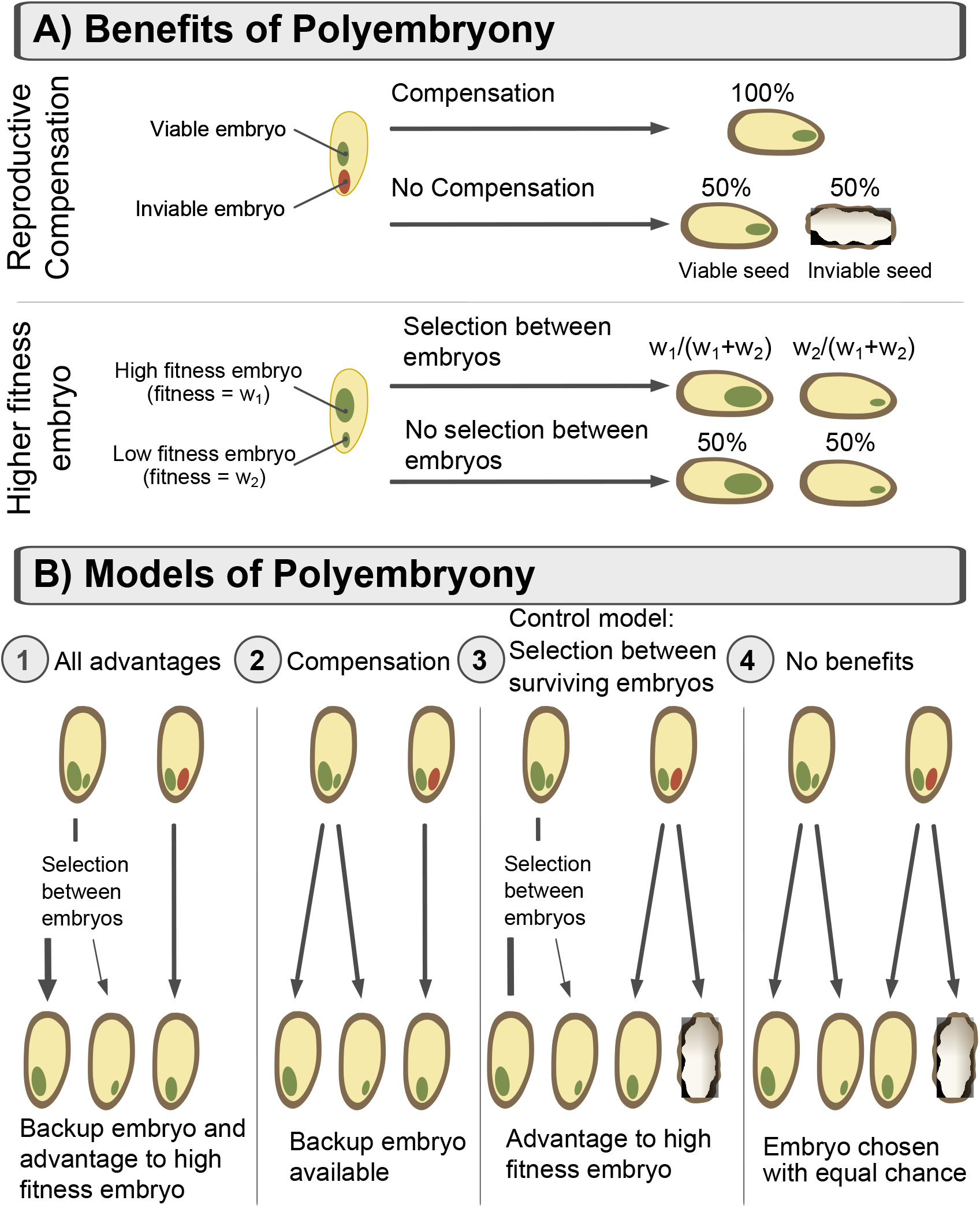
Benefits ad models of polyembryony: (**A**) Polyembryony offers two potential benefits – increasing the chances having a viable seed (Reproductive compensation, top panel), and increasing the chances that a high fitness embryo is in the seed (Selection between surviving embryos, bottom panel). (**B**) Factorially combining these benefits generates our four models for the evolution of polyembryony (see text for details).

The reproductive compensation benefit of polyembryony is that having two potential embryos in a seed increases the probability that a seed will contain a viable embryo (Top panel of Figure 2A). Thus, the benefit of reproductive compensation occurs if the survival of at least one embryo ensures a viable seed. Conceptually and biologically, it is somewhat difficult to remove the benefit of reproductive compensation while maintaining the benefit of a getting higher fitness embryo, because higher fitness embryos are more likely to survive than low fitness embryos. Therefore some reproductive compensation will naturally accompany selection between embryos. Although these benefits are difficult to dissect biologically, we can dissect them computationally by introducing a control variant of our model in which we limit selection between embryos to cases in which both embryos survive. This ’selection among surviving embryos’ control effectively removes the benefits of reproductive compensation by allowing seed viability to be determined by survival of an arbitrarily chosen embryo among that seed’s two embryos. We note however, that this control does not line up clearly with simple biological scenarios and is therefore presented to provide conceptual clarity, not biological realism.

Factorially combining these options results in four models for the evolution of polyembryony (Figure 2B). As we describe these models, we show how each model isolates the potential outcomes of polyembryony by considering the survival of focal embryo one, *p*_1_, in a seed with a second embryo. We assume embryos one and two have survival probabilities *w*_1_ and *w*_2_, respectively, and we note the probability that embryo one is chosen by embryo screening as 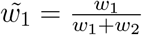.

#### 1. All advantages

In this model, each embryo in the seed of a polyembryonic mother survives independently with a probability determined by its embryonic fitness. If only one embryo survives, this embryo develops into the seed, and if both embryos survive, the embryo that develops in the seed is chosen in proportion to the relative embryonic fitness of each embryo (i.e. if both embryos survive, the probability an embryo develops into a seed equals its fitness divided by the sum of the fitness of both surviving embryos). Thus, this model includes both potential benefits of polyembryony.

In the all benefits model, the probability that embryo makes it to the seed is the probability it survives and embryo two dies (*w*_1_(1 *− w*_2_)) plus the probability that both survive, and embryo one ends up in the seed 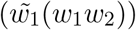.

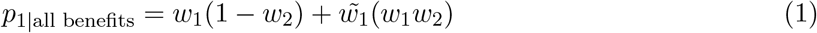

#### 2. Compensation only

In this model, each ovule in the seed of a polyembryonic genotype survives independently with a probability determined by its embryonic fitness. If only one embryo survives, this embryo develops into the seed, and if both embryos survive, the embryo that develops in the seed is chosen at random. As such, polyembryony provides the benefit of increasing the probability that a seed survives, but does not provide the added benefit of favoring the higher fitness embryo over the lower fitness one. This model resembles the case of reproductive compensation (Porcher and Lande 2005), as inviable genotypes can be replaced. With this type of selection, embryo viability is not dependent on the fitness of the other embryo, however embryos with inviable siblings are more likely to become seed than are embryos with viable siblings.

Thus, in the compensation model, the probability that embryo makes it to the seed is the probability it survives and embryo two dies (*w*_1_(1 *− w*_2_)) plus the probability that both survived and that this embryo was randomly chosen to end up in the seed (*w*_1_*w*_2_/2).

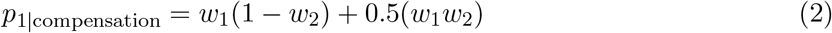

This equation is similar to the all benefits model (Eq. 1), with the difference that 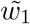 is replaced by 0.5 highlighting the absence of slection between surviving embryos in the compensation only model.

#### 3. Control model: No reproductive compensation

We can indirectly infer the advantages and consequences of selection between surviving embryos by comparing the all benefits and compensation models above. However, directly modelling the benefits and consequences of allowing for selection between embryos without allowing for reproductive compensation is somewhat complex for two reasons. The first challenge is that if an embryo’s survival probability is directly related to its relative fitness, than allowing for selection between embryos includes the benefit of reproductive compensation. Second, if embryo viability is unrelated to survival probability it is difficult to model the evolution of relative embryo fitness when there is only one embryo in a seed (i.e. before the introduction of polyembryony). We therefore developed the following approach to model allow for embryo selection selection before and after polyembryony evolved while removing the natural advantage of reproductive compensation associated with polyembryony. In this model, a seed in a polyembryonic genotype survives with a probability equal to the embryonic fitness of an arbitrarily chosen embryo (embryo 1 in our simulation). Thus, if this embryo dies but the other lives, the seed still dies. As such, monoembryonic and polyembryonic genotypes will have the same probability of developing a viable seed (i.e. in both cases seed survival is determined by the fitness of a single random embryo). However, if both embryos survive, the embryo that develops in seed is chosen in proportion to the relative embryonic fitness of each embryo. This control variant maintains the benefit of selection between embryos, while removing the benefit of reproductive isolation.

Thus, in the model of ’selection among surviving embryos’ without reproductive compensation, the probability that embryo makes it to the seed is the probability it survives and embryo two dies divide by two (0.5(*w*_1_(1 − *w*_2_))) plus the probability that both survive, and embryo one ends up in the seed 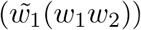.

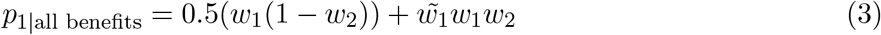

We note that this outcome is similar to the all benefits model (Eq. 1), with the difference that *w*_1_(1 *− w*_2_) is weighted by 0.5, so as to effectively removes the benefit of reproductive compensation from this model, while maintaining selection between surviving embryos.

However, while our model of reproductive compensation without selection between embryos seems biologically plausible, our model of selection between surviving embryos without reproductive compensation does not – it is unclear how survival of a seed could be determined by the fitness of a random embryonic genotype it carries that may or may not end up in the seed. As such, this scenario is introduced, not as an attempt to model biological reality, but rather to provide a theoretical control to allow us to better understand the biological world. That is, while biology may not allow for selection between embryos without an incidental benefit of enhanced embryo survival our modeling approach can, which allows us to isolate the evolutionary consequences of each.

#### 4. No benefit

In this model, one of the two embryos in the seeds is randomly chosen to make it into the seed, and the seed is inviable if this embryo is inviable. As such, there should be no advantage of polyembryony. This model acts as a control to ensure that our simulation scheme meets neutral expectations and that our control for each potential benefit of polyembryony is properly implemented.

## Results

### Burn-in and the evolution of the load

We discuss the results from our burn-in simulations, as they set the scene for the evolution of polyembryony. Throughout the discussion of burn-in results, we focus on mutations impacting embryo fitness, as results for post-embryonic fitness follow similar qualitative and quantitative patterns (Figure S1). Genomes saved at the end of the burn-in are available for download here.

### Comparison to published analytical results

Before discussing specific results, we evaluate whether our simulations behave sensibly by comparing model output to known analytical results – namely, the expected number of recessive lethal mutations per diploid genome in a panmictic population. Based on classic results of Li and Nei (1972), Gao et al. (2015) show that the expected number of recessive lethals per (diploid) individual in a finite, panmictic population equals 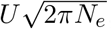, where *U* is the mutation rate per haploid genome, and *N*_*e*_ is the effective population size. For the case of recessive embryo lethals in outcrossers, we find a mean of 18.6 mutations per diploid genome, a value remarkably consistent with the predicted value of 18.4 (Compare dashed white line to simulation results in Figure 3A). That is, if *U* = 0.25, as we are only concerned with mutations impacting embryo fitness (half of total mutations), and 2*N*_*e*_ *≈* 1745, the mean number of surviving embryos across replicates in the final generation. Additional exploratory simulations (not shown) found a consistent agreement with theory across a range of mutation rates.

**Figure 3:**
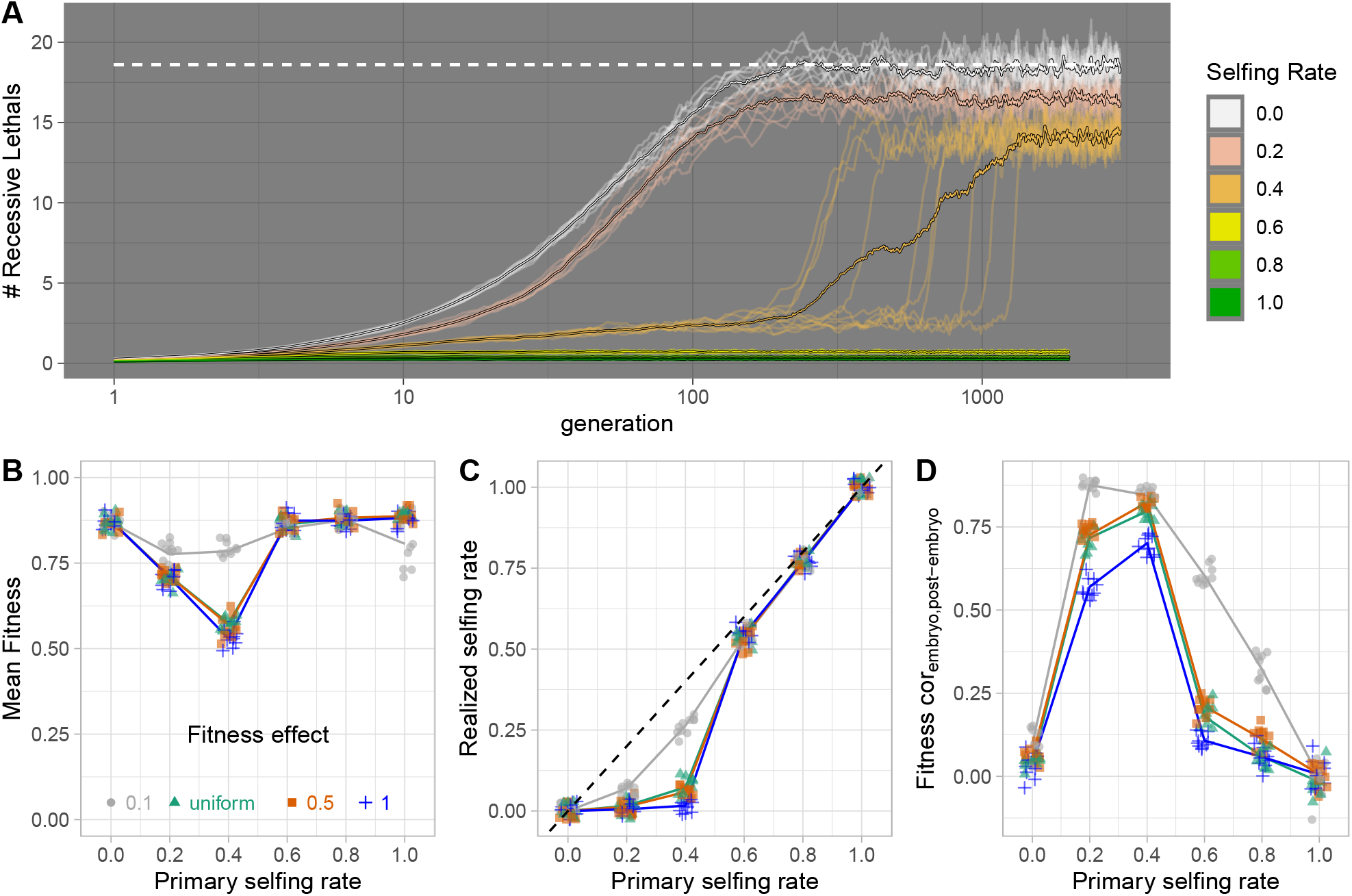
Results from burn-in: **(A)** The mean number of recessive lethal alleles per haploid genome over time. Each line is one of ten replicates for each selfing rate, designated by color. The dashed white line shows the theoretical expectation for a randomly mating population in our simulation, while the larger colored line shows the mean across replicates. Results with different dominance and selection coefficients are presented in Figure S1. Figures **(B-D)** show features of the population ’burn-in’ populations after the load equilibrates. Points are slightly jittered to show the data - with one value for each replicate simulation for a given combination of selfing rates on the x, and fitness effects of new mutations in color, lines connect means. In **C**, the one to one line is shown by the dashed black line. All mutations are fully recessive. Results with different dominance coefficients are presented in Figure S2.

### Novel Burn-in Results

Recessive lethal mutations are effectively purged with predominant selfing (selfing rate > 0.5), while a large number of deleterious mutations accumulate with predominant outcrossing (Figure 3A). With an intermediate selfing rate of 0.4, the population appears to reach an equilibrium, relatively modest number of recessive mutations, until this rapidly and dramatically increases, presumably reflecting a transition from effective purging to interference among deleterious mutations (Lande et al. 1994; Porcher and Lande 2016). Across all parameter combinations, the number of deleterious mutations at equilibrium decreased as mutations became more deleterious and more additive (Figure S1A). Additionally, across all dominance and selection coefficients, the number of deleterious alleles in a population decreased with the selfing rate. However, the results for obligate selfers were somewhat unstable with weak selection and non-recessive dominance coefficients, when mutations can occasionally fix, and therefore continue increasing in number (Figure S1).

When mutations are recessive (*h* = 0), mean fitness is lowest with intermediate selfing rates, and is generally highest with high levels of selfing or outcrossing (Figure 3B). This pattern is most pronounced when recessive mutations are lethal (*s* = 1), exceptionally deleterious (*s* = 0.5), or where selection coefficients were drawn from a uniform distribution, and more subtle with a selection coefficient of *s* = 0.1 (Figure 3B). By contrast, when mutational effects are additive (*h* = 0.5) or are drawn from a uniform distribution, mean fitness increases with the selfing rate, with significantly positive slopes ranging between 0.045, and 0.082 (Figure S2A, Table S1), presumably because selfing increases the variance in fitness, allowing for more effective selection. In these non-recessive cases, mean embryo fitness is roughly similar, regardless of the fitness effects of individual mutations (Figure S2, modelling mean fitness = *f* (selfing, *s*), the p-value for the effect of *s* is 0.059 and 0.25, for cases with a uniform and additive load, respectively, Table S2). Reassuringly, this grand mean fitness under obligate outcrossing for non-recessive alleles of 0.78 is in line with its predicted value of 0.78 from Haldane’s 1937 classic result that mean fitness equals *e*^*−U*^ (where *U* is the mutation rate per haploid genome, which equals 0.5 divided by two, as half of mutations will impact embryo fitness). Somewhat surprisingly, mean post-embryonic fitness does depend on the selection coefficient (Fig. S2D), suggesting that selection at one life stage impacts outcomes at another as suggested by Sakai (2019).

With intermediate selfing rates and recessive gene action we observe a much higher primary than realized selfing rate, suggesting that inbreeding depression underlies much of the embryo death in these cases (Fig. 3C). By contrast, we observe a nearly perfect relationship between primary and realized selfing rates under non-recessivity (Fig. S2B). We observe a strong positive correlation between embryo and post-embryo fitness for recessive gene action and intermediate selfing rates, but no relationship otherwise (Fig. 3D, and Fig. S2C). Together these results support the intuition that if competition acts to remove selfed embryos, this benefit of polyembryony will be most relevant when mutations are recessive.

### Invasion of polyembryony

We compare the fixation probability of a new mutant that confers polyembryony, across all models described above. We find that, when the polyembryony allele fixes, it tends to fix more quickly when polyembryony provides reproductive compensation than when it does not (Fig. 4A, Fig. S4, Table S3). Similarly, polyembryony is most likely to fix when it provides reproductive compensation – in some cases, single mutations have up to a fifteen percent chance of reaching fixation, a 300-fold increase in the probability, relative to neutral expectations (Fig. 4B & 4C). Our control model, which removed the benefit of reproductive compensation but maintained the benefit of selection between surviving embryos, also favored the evolution of polyembryony, but had a more modest effect – in some cases, single mutations have up to a one percent chance of reaching fixation, a 20-fold increase in the probability, relative to neutral expectations. Reassuringly, fixation proportions from the no benefits model matched neutral expectations, with approximately 1/2*N* = 0.0005 introductions resulting in fixation (See Table S3, and compare the solid lines to the dashed line in Figure 4B).

**Figure 4:**
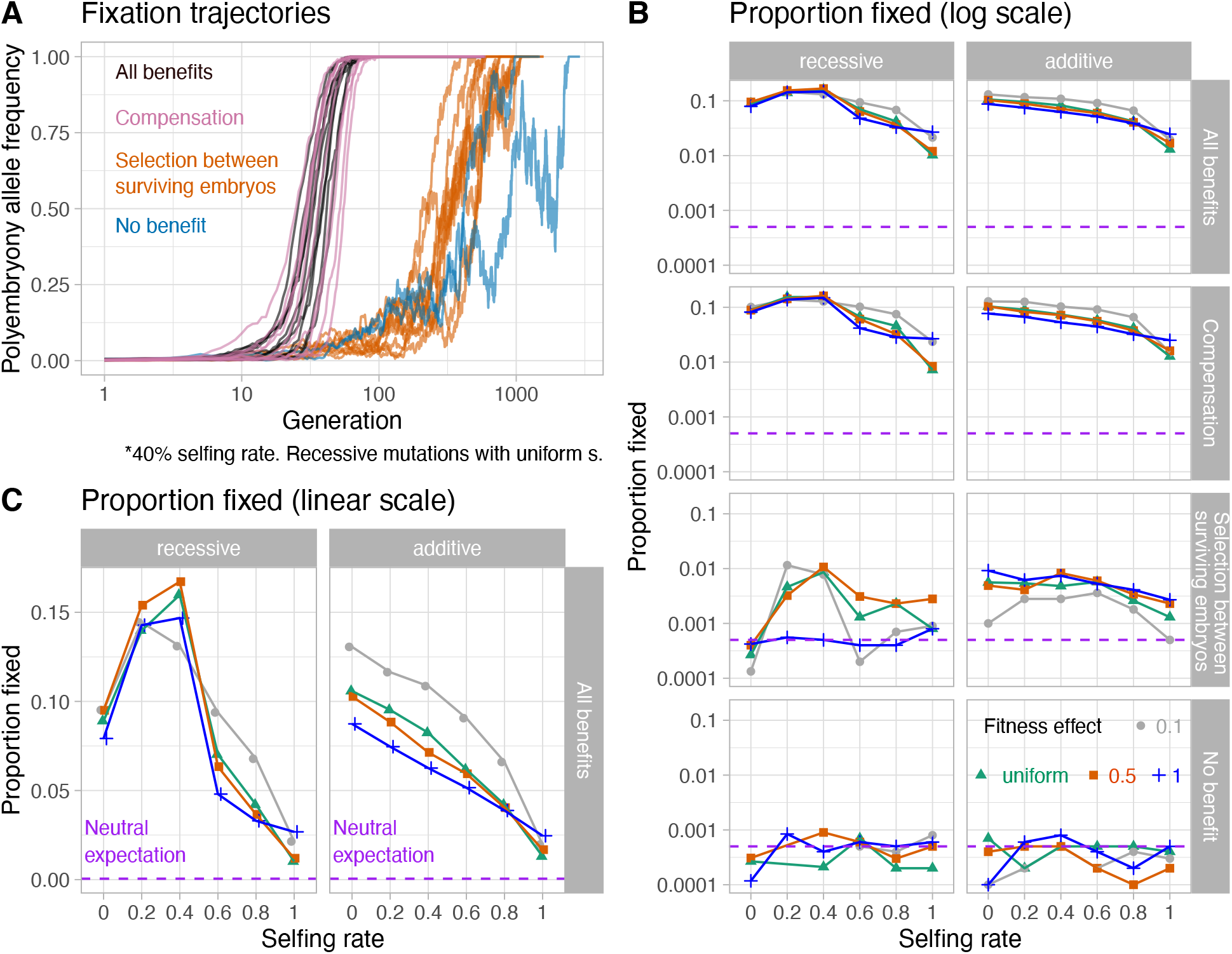
The fixation of an allele conferring polyembryony: **A)** Example trajectories of the fixation of the polyembryony allele with all benefits (black), the benefit of compensation (pink), the benefit of selection between surviving embryos without reproductive compensation (orange), and no benefit (blue). Note that generation on x increases on the log10 scale, but specific values are noted with their linear value. **B)** The proportion of introductions resulting in fixation of the polyembryony allele as a function of the selfing rate (x), the fitness effect of new mutations (color), the mode of gene action (columns), and the benefit of polyembryony (rows). The dashed pink line displays the expectation under neutrality. Note that fixation proportion on y increases on the *log*_10_ scale, but specific values are noted with their linear value. **C)** The proportion of introductions resulting in fixation of the polyembryony allele as a function of the selfing rate (x), the fitness effect of new mutations (color), the mode of gene action (columns), with all the benefits of polyembryony. The values are identical to those in the first row of **B**, but are presented on a linear scale to highlight the effect of selfing rate on fixation probability.

Other biological parameters such as the selfing rate, and the dominance and selective coefficients of deleterious mutations also impact on the evolution of polyembryony, often depending on their interaction. Below we discuss the effects of selfing rate and additive vs. recessive modes of gene action, noting that results from the uniform mode of gene action are qualitatively similar to the additive model (Fig. S3, Table S3).

#### The benefit of reproductive compensation

strongly favored the evolution of polyembryony for all biological parameters investigated (Fig. 4). Figure 4C displays the fixation proportions for the compensation models (row two in Fig. 4B) on a linear scale to reveal the effect of selfing rate and selective effects of new mutations.

Under recessivity, the probability of fixation is maximized (approximately 15%) at intermediate selfing rates, suggesting that polyembryony can evolve to make up for offspring lost to early acting-inbreeding depression. Again assuming recessivity, obligate outcrossing more strongly favors the evolution of polyembryony than does obligate selfing (compare an approximately 10% fixation probability under random mating to a 2.5% fixation probability under obligate selfing, Fig. 4C), presumably reflecting the higher within-seed variance in fitness under obligate outcrossing leading to higher impact of polyembryony. The fitness effect of recessive deleterious mutations have only a modest effect on fixation proportion, varying slightly across selfing rate.

However, the compensation model also strongly favors the evolution of polyembryony with an additive load, suggesting that overcoming inbreeding depression is not the only driver of the evolution of polyembryony. (Second row, second column, Figure 4B). In cases with additive gene action, the fixation probability of a polyembryony allele decreases with the selfing rate, again reflecting the lack of within-seed variance in fitness (i.e. the benefits of replacing an inviable embryo decrease as the probability that its replacement will also die increases). Additionally, under additivity (or if mutations take their dominance coefficients from a uniform distribution, Table S3, Figure S3) a load composed of highly deleterious mutations is less likely to foster the evolution of polyembryony than a load composed of a larger number of mild mutations (compare *s* = 1 (blue) to *s* = 0.5 (orange) or s=uniform (teal) to *s* = 0.1 (grey), Fig. 4). This surprising result might reflect the fact that while mean fitness does not depend on fitness effects of new mutations, the survival of maternal sibling-embryos becomes more dependent on one another as mutational effects get larger (Figure S5). As such, with large effect mutations, a backup embryo is less useful as if one dies the other is likely to die as well.

#### Removing the benefit of reproductive compensation, while allowing for selection between surviving embryos

also favors the evolution of polyembryony. However, fixation probabilities are approximately five-to ten-fold lower for this model than for the reproductive compensation model. With a recessive load and intermediate selfing rates (0.20 or 0.40), this model results in the fixation of the polyembryony allele in approximately one percent of introductions, a twenty-fold increase relative to the neutral expectation of 0.05%. Somewhat surprisingly, selection between surviving embryos favors polyembryony for a non-recessive load (Third row, second column of Fig. 4B), even though embryo fitness was uncorrelated with post-embryo fitness in these models (Fig. S2). This likely reflects the benefit of producing grand-children with higher embryonic fitness who will out-compete their siblings (analogous to models of “runaway sexual selection” Kirkpatrick 1982). Under both additivity and a uniform distribution of dominance effects, the probability of fixation of a polyembryony allele that does not allow for reproductive compensation but allows for selection between surviving embryos is greatest in predominantly outcrossing populations (selfing rates of 0.40 or less), decreasing as the selfing rate increases. Here, the probability of fixation is greatest when the load is composed of alleles of large effect, a result that runs counter to that found in the compensation model with an additive load.

#### All benefits

results in fixation probabilities qualitatively similar to the reproductive compensation model (Fig. S3, Table S3) – reflecting the importance of the benefits of reproductive compensation, to the evolution of polyembryony.

#### No benefits

results in fixation probabilities consistent with neutral expectations (Fig. S3, Table S3).

### Evolutionary consequences of polyembryony

We compare how different models of the evolution of polyembryony shape key features of a population, including the proportion of surviving seeds, the realized selfing rate and the architecture of genetic load. Although a strict version of our control model which removes the benefit of reproductive compensation but allows for selection between surviving embryos is unlikely to occur in nature, its inclusion allows us to distinguish the individual effects of these two benefits when both would be operating in nature (i.e. the all benefits model). Because results were qualitatively similar across all selection coefficients (save the decrease in fitness with recessive mutations, *s* = 0.1 and high selfing rates, which did not always converge Fig. S1), and because results from the additive model and the uniformly distributed dominance coefficient model did not differ qualitatively, we focus on results from the cases in which the selection coefficients of new mutations are selected at random from a uniform distribution, exploring cases in which mutations are recessive or additive (Figure 5).

**Figure 5:**
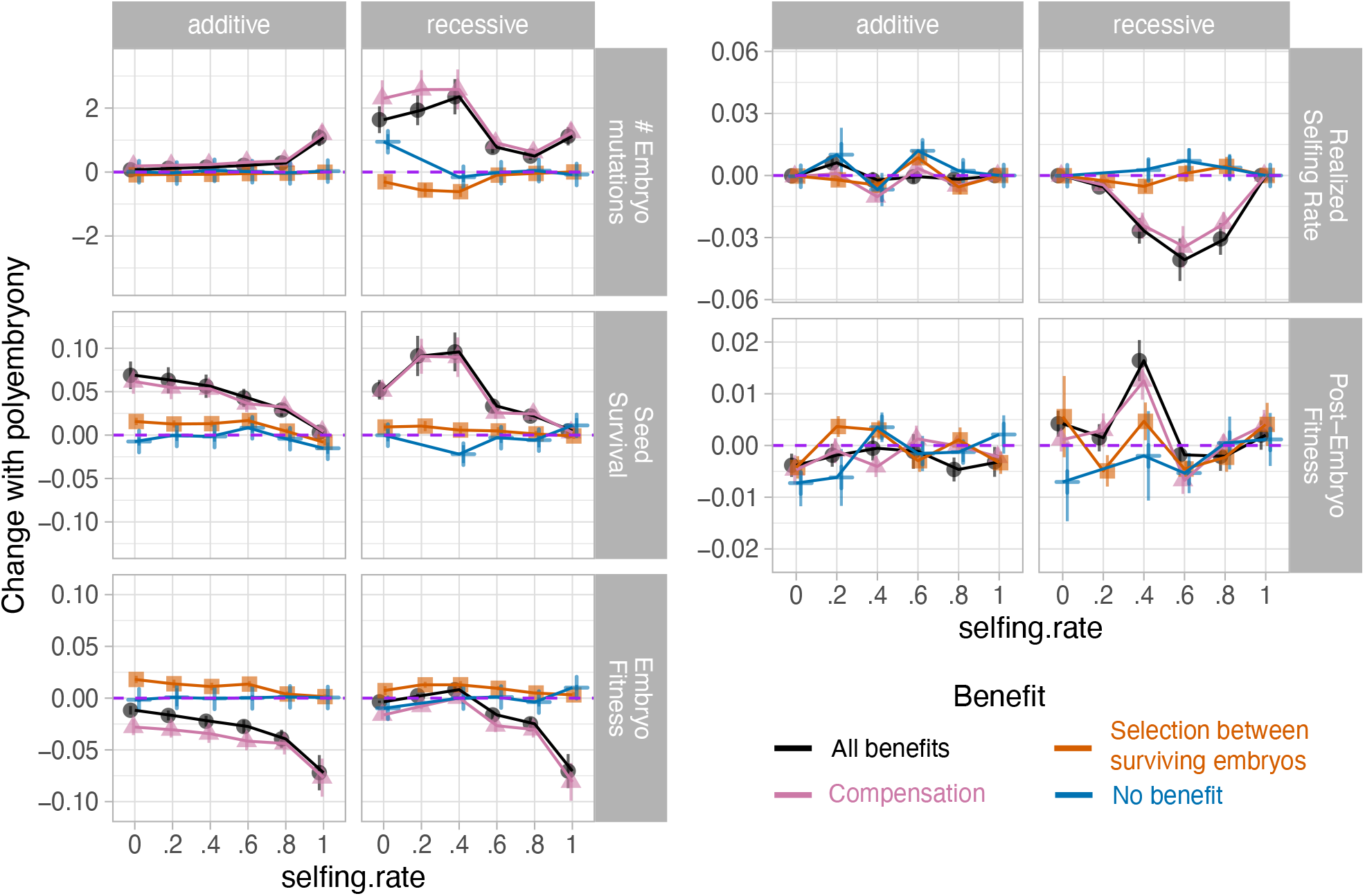
The evolutionary impact of polyembryony. How the evolution of polyembryony impacts the per individual number of mutations impacting embryo fitness, expected seed survival, mean embryo fitness, the realized selfing rate, and mean post-embryo fitness (rows), when mutations are additive or recessive (columns), across selfing rates (x-axis), for each model of polyembryony (color).

Curiously, the allowing for selection between surviving embryos but not reproductive compensation did not impact the realized selfing rate (Fig. 5), even with recessive mutations and intermediate selfing rates. This result is a consequence of features of both our model and biological reality. Specifically, there are limited opportunities for selection between surviving selfed and outcrossed embryos in a seed (Williams 2007), as this only occurs with a probability equal to two times the variance in the selfing rate (the probability that exactly one of two embryos is from a self-fertilization event) times the probability that both are destined to survive. By contrast, with a recessive load a benefit of compensation decreases the realized selfing rate, and increases the number of mutations impacting embryo fitness in partially selfing populations (Fig. 5).

The benefit of reproductive compensation (in both the compensation and all benefits model) increased seed survival. Under a recessive load, this effect was maximized with intermediate selfing rates, while it decreased steadily with selfing rate under a (partially) additive load. Allowing for selection between surviving embryos without reproductive compensation subtly increased seed survival for all models of dominance investigated so long as the selfing rate was not too large (Fig. 5). Consequently, the expected embryo fitness of the surviving seeds subtly increases with the benefit of selection between surviving embryos, but decreases with compensation. These consequences are additive, such that the expected embryo fitness decreases in the all benefits model but does so less severely than in the compensation model.

Regardless of the mode of gene action, selection between surviving embryos without reproductive compensation does not increase the expected post-embryo fitness of surviving seeds (Fig. 5). Together, these lines of evidence suggest that polyembryony does not evolve as a mechanism to prevent self-fertilization, and is not analogous to the system of self-incompatibility observed in angiosperms.

Our models show that the benefit of reproductive compensation favors the evolution of polyembryony. Could the availability of a ’back-up’ embryo encourage the evolution of self-sabotage of selfed seeds as has been suggested as an explanation for the embryonic lethal system? According to this model, polyembryony evolves as a mechanism of reproductive compensation, but the embryolethal system evolves as an SI-like mechanism to destroy selfed seeds. Figure 6 shows that the allele frequency spectrum is comparable in the no benefit and the selection between surviving embryo models, arguing against the idea that selection between surviving embryos favored self-sacrifice in the form of an excess of rare recessive lethals. By contrast, there is a slight increase in the count of deleterious recessive mutations across all frequency classes in the compensation and all benefits models. It is unclear it this shift reflects the relaxation of embryo selection in these cases or selection favoring self destruction of selfed embryos.

**Figure 6:**
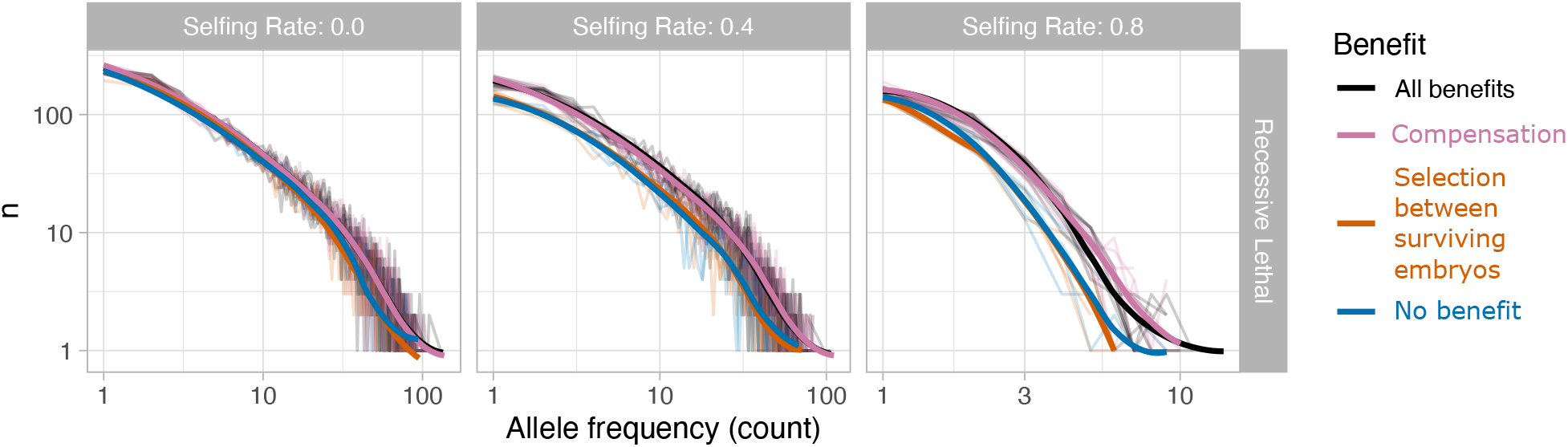
The allele frequency spectrum for embryo acting allele following the evolution of polyembryony under the recessive lethal model for a selfing rate of 0.00, 0.40, and 0.80 (left to right). Lines display averages of ten simulation replicates, and colors note the model of polyembryony. Note that the x axis with a selfing rate of 0.80 (right panel) is truncated relative to the other selfing rates, reflecting the effective purging of early acting recessive mutations with high selfing rates.

## Discussion

We present four models to test the plausibility of the compensation and ”fitter offspring” theories for the evolution of polyembryony. We find that the evolutionary benefit of compensation – that is, the opportunity for a backup embryo to replace an inviable one – strongly favors the evolution of polyembryony. Relative to neutral expectations, the benefit of compensation results in between a twenty-fold increase in fixation probability above the neutral expectation with high selfing rates, and a two hundred-fold increase with intermediate to low selfing rates and a recessive load, all across a broad range of selection and dominance coefficients. By contrast, removing this benefit while allowing for selection between surviving embryos more weakly favored the evolution of polyembryony, resulting in between a zero-fold increase with high selfing rates, and a twenty-fold increase, with intermediate to low selfing rates and a recessive load, relative to neutral expectations. As such with both benefits, fixation probabilities closely follow expectations from the compensation model alone.

Our work highlights the previously underappreciated result that the consequences of polyembryony depend on its function. When polyembryony functions as a compensation mechanism, mean embryo fitness is reduced, but the probability of seed survival increases, consistent with previous work (e.g. Klekowski 1982; Porcher and Lande 2005). By contrast, when competition between embryos acts only to facilitate selection between surviving embryos, mean embryo fitness increases, seed survival does not change substantially, in line with models of competition alone (Latta 1995). With a recessive load, the benefit of compensation acts to decrease the effective selfing rate, while the selection among surviving embryos did not. With both the benefits the evolutionary consequences of polyembryony is somewhere in between but is often closer to those expected from compensation.

### The limited role of selection among viable embryos in the evolution of polyembryony

It has long been assumed that a major benefit of polyembryony is that it provides an opportunity for embryos to compete (Sarvas 1962; Koski 1971), and therefore for mothers to screen for high fitness offspring. Not only was the benefit of allowing for selection between surviving embryos itself – after removing the potential benefit of reproductive compensation – a comparatively weak force in the evolution of polyembryony, but it did not reliably increase seed fitness. In fact, under most selfing rates and dominance coefficients, selection between surviving embryos more strongly favored the evolution of polyembryony with an additive load (in which there is no relationship between embryo and post embryo fitness) than a recessive load (in which there was such a relationship). This is surprising because there is a limited scope for selection on polyembryony when it cannot affect post-embryo fitness. In this case, selection on polyembryony only occurs within polyembryonic seeds themselves, and, since the embryos’ maternal genomes are identical, only among the paternal genomes. Like runaway sexual selection (Fisher 1915), the automatic transmission advantage of selfing (Fisher 1941), or meiotic drive (Rhoades 1942), selection between surviving embryos is an example of a selective advantage that does not make a population necessarily more adapted to its environment.

### Why doesn’t selection between surviving embryos effectively remove selfed offspring and increase post-embryonic fitness (and could it ever)

In our control model of selection between embryos without reproductive compensation, between embryo selection is only relevant if both embryos were destined to live. As such, this mechanism is most effective at removing selfed offspring when one seed is selfed and the other is not, and both seeds are destined to live (because if, for example, the selfed embryo was destined to die, reproductive compensation would allow for the replacement of this low-fitness, selfed embryo). Because predominant selfing (selfing rate > 0.5) purges the recessive load, and predominant outcrossing (selfing rate *<* 0.5) generates large inbreeding depression, in most cases in which one embryo is selfed and the other is outcrossed, their fitness is either nearly equal, or the selfed embryo is destined to die. As such, selection between surviving embryos does not offer a more refined view into post-embryo fitness than is automatically accounted for by “hard selection” on seed viability. Our observation that selection between surviving embryos leads to more competitive embryos, rather than higher fitness plants is consistent with the claim of McCoy and Haig (2020) that Goodhart’s law – ‘When a measure becomes a target it ceases to be a good measure’ – can undermine effective embryo selection.

Despite our focus on the evolution of polyembryony, these results apply broadly and suggest that verbal models predicting that selective embryo abortion could limit the mating costs of selfing in plants with mixed mating systems (e.g. Huang et al. 2020), require more rigorous scrutiny.

Nonetheless, it is possible that pure ”soft selection” (Wallace 1968, 1975) on embryos could reliably increase post-embryo fitness. That is, although we had trouble implementing this model computationally (e.g. we could not define the allele frequency spectrum of alleles determining success in soft selection before embryo screening evolved), nor could we map this onto a plausible biological mechanism, these challenges could reflect a shortcoming in our imagination, rather than a biological impossibility. Additionally, we note that even if the benefits of compensation initially favored the evolution of polyembryony, it is possible that the evolution of polyembryony was followed by novel recessive mutations experiencing soft selection, and therefore the benefits of between embryo selection could represent an exaptation (Gould and Vrba 1982) maintain but not drive the evolution of polyembryony.

### The embryo lethal system

Since Buchholz (1922), it has been argued that the embryo-lethal system, an apparently coordinated process of embryo death, could achieve a similar function to angiosperm self-incompatibility in the self-compatible gymnosperms. That is, coordinated death in the embryo stage would give way to highly outbred surviving adults (Sarvas 1962; Koski 1971). This would be an altruistic act in which an embryo sacrifices its predictably low fitness for a maternal half sibling. We did not observe the evolution of an embryo lethal system in response to the evolution of polyembryony, as would be expected if polyembryony favored altruistic self-destruction of more inbred embryos (e.g. we did not see a change in the allele frequency spectrum towards an excess of low-frequency recessive mutations). However, by relaxing selection on embryo viability, embryo compensation could indirectly result in an increase in the number of highly deleterious recessive mutations impacting embryo fitness. As such it appears that the embryo lethal system could reflect an elevated load tolerance rather than an exquisite adaptation (Gould and Lewontin 1979), as Williams (2007) argued forcefully based on developmental and genetic evidence. However, differentiating between these alternatives require additional modelling efforts.

Finally, we note that we cannot exclude possibilities which we did not model (e.g. pure soft selection). For example, in a single locus, 4 allele haploid system with pure soft selection, Archetti (2009) found that an “anti-robust” allele that amplified the deleterious effects of a mutation in juveniles so as to be replaced by a genotype that will result in a higher fitness adult will be favored. While one could interpret the “embryo lethal system” as an anti-robust system, it is unclear if such alleles could evolve in a complex diploid genome (e.g. pines).

### Which has driven the evolution of polyembryony – Compensation or selection between embryos

We find that the benefit of embryo compensation favors the evolution of polyembryony more strongly than does selection between surviving embryos. However, we caution that whether compensation or selection between embryos have actually favored the evolution of polyembryony depends on their biological plausibility and whether they reflect effective solutions to the problems they address. That is, we must consider biological processes outside of our model as we interpret our model results. For example, favoring a fitter embryo could perhaps be most effectively achieved by placing more embryos in a seed, while compensation could be more effectively achieved by producing more seeds per plant. Throughout this discussion, we remind the reader that our model of selection between surviving embryos was not meant to model biological reality, but should provide insight into how selection between embryos could influence the evolution of polyembryony and its consequences – allowing us to infer which benefit most strongly contributes to the evolution of polyembryony in nature (where these two features cannot be clearly disentangled).

Our models provide competing testable predictions to distinguish between predictions of the compensation and the fitter embryo model at within seed level, for simple polyembryony, assuming no pleiotropic effects. For example, we show that the evolution of polyembryony and its consequences depend on the selfing-rate and dominance coefficient. Specifically, with a recessive genetic load, the opportunity for selection between surviving embryos most strongly favors the evolution of polyembryony at intermediate selfing rates (Fig. 4B,C). The estimates of selfing rates for modern conifers can reach 0.30 - 0.60 (Sarvas 1962; Sorensen 1982), a range that favors polyembryony. We note, of course, that estimates of the primary selfing rate from extant conifers rate may differ substantially from the primary selfing rates of the population in which polyembryony arose.

Additionally, the two models make subtly different predictions about the difference between the realized and primary selfing rate. Relative to a monoembryonic ancestral population, polyembryony favored by the opportunity for selection between embryos but not reproductive compensation does not result in decrease in the difference between realized and primary selfing rates. By contrast, with a recessive load and intermediate selfing rates, polyembryony favored by compensation strongly amplified the difference between the realized and primary selfing rates. In nature, differences between primary and realized selfing rates are often observed in species with simple polyembryony (Lindgren 1975; Sorensen 1982; Kärkkäinen and Savolainen 1993; Lande et al. 1994), further emphasizing the probable role of compensation in the evolution and maintenance of polyembryony.

### The impact of pleiotropy across the life cycle

In the main results presented above, We assumed no pleiotropy across life stages – that is, mutations either impacted embryo or post embryo fitness. However, this is clearly untrue in nature. For example, severe loss of function mutations in key genes would likely decrease both embryo and post-embryonic fitness. We therefore wanted to evaluate the extent to which our results rested on the assumption of no pleiotropy. To do so, we re-ran all models under the assumption of complete pleiotropy – in which all mutations had identical fitness effects on embryonic and post-embryonic fitness.

Somewhat surprisingly, a model of exceptionally strong pleiotropy – in which each mutation had an identical effect on both embryonic and post-embryonic fitness – showed that reproductive compensation strongly favored the evolution of polyembryony, while selection between surviving embryos less reliably resulted in the fixation of polyembryony (Figure S6. Together, these results suggest that embryo compensation is likely the major selective force favoring the evolution of polyembryony,

We did not explore the case in which an allele antagonistically increased embryo fitness while decreasing post embryonic fitness. When such a mutation occurs with embryo screening, it could generate an ontogenic conflict. Empirical studies, e.g. mapping and measuring of inbreeding depression at different life stages (Koelewijn 1998), comparing gene expression across embryo development and later life stages (Raherison et al. 2012), and signatures of negative and positive selection in such genes would be valuable to further evaluate the potential importance of pleiotropy in the evolution of polyembryony. However, as our results suggest that reproductive compensation is the major driver of the evolution of polyembryony, it seems like this ontogenic conflict would play only a minor role in its evolution.

Finally, since polyembryony is primarily favored by the benefit of reproductive compensation, this benefit should be independent of the mechanism underlying embryo inviability. As such, polyembryony may be favored as a form of reproductive compensation for embryos whose viability is compromised by developmental hazards unrelated to genotype.

### Selection between embryos, compensation, and conflict in a pine nutshell

Gymnosperm seed with a maternal haploid megagametophyte, multiple genetically distinct embryos, genetically identical (cleavage) embryos, and strong inbreeding depression is a stage of evolutionary drama that deserves more attention, and we hope that the provided model will be used to broaden the investigations on the evolutionary dynamics outside the angiosperm sphere. For example, in contrast to the opportunity for altruism to favor the embryo-lethal system, polyembryony also provides avenues for parental and embryonic conflict.

In simple polyembryony, embryos are potentially derived from different sires. A paternal genome carrying a mutation that sabotaged rival embryos carrying different paternal genomes could possess a net advantage even if doing so would reduce the probability that a viable seed is formed at all. Sabotage and anti-sabotage alleles would only be beneficial when expressed in a particular parental genome, so genomic imprinting that prevented expression in the wrong parental genome would also be advantageous.

Conifers and other gymnosperms provide unique opportunities to test key questions of plant mating system evolution and evolutionary conflict from a novel angle, especially now that their genomic resources are no longer seriously hindered by their large genome sizes (e.g. Niu et al. 2022). From the empirical perspective, large seed size and gametophytic tissue allow easy identification of maternal haplotypes and alleles. Thus expression patterns and genetic diversity for example in potentially imprinted genes should be easy to quantify and identify in many conifer species.

## Supplement

**Table S1:** Slope of the relationship between selfing rate and mean embryo fitness after burn-ins for non-recessive variants. All t values are associated with 59 degrees of freedom.

| **h** | **s** | **estimate** | **lower 95% CI** | **upper 95% CI** | **$t$** | **p-value** |
| --- | --- | --- | --- | --- | --- | --- |
| 0.5 | 0.1 | 0.054 | 0.031 | 0.076 | 4.719 | 0.000015 |
| 0.5 | 0.5 | 0.085 | 0.076 | 0.094 | 19.114 | <10-11 |
| 0.5 | 1 | 0.051 | 0.039 | 0.062 | 8.600 | <10-11 |
| 0.5 | uniform | 0.082 | 0.072 | 0.091 | 16.413 | <10-11 |
| uniform | 0.1 | 0.045 | 0.015 | 0.074 | 2.980 | 0.004203 |
| uniform | 0.5 | 0.078 | 0.068 | 0.088 | 15.313 | <10-11 |
| uniform | 1 | 0.065 | 0.055 | 0.076 | 12.021 | <10-11 |
| uniform | uniform | 0.083 | 0.074 | 0.091 | 19.445 | <10-11 |

**Table S2:** Effect of selection coefficient on mean fitness following burn-in for non-recessive variants.

| **h** | **estimate** | **$F_{3,225}$** | **p-value** |
| --- | --- | --- | --- |
| 0.5 | 0.054 | 1.39 | 0.059 |
| uniform | 0.083 | 2.52 | 0.247 |

**Table S3:** The proportion of introductions of the polyembryony allele resulting in fixation (No compensation refers to the control model with selection among surviving embryos, which provides the benefit of a fitter embryo but precludes reproductive compensation).

| $p_{\text{self}}$ | **s** | **h** | **All benefits** | **Compensation** | **No compensation** | **No benefit** |
| --- | --- | --- | --- | --- | --- | --- |
| 0 | uniform | uniform | 0.1084 (n=5000) | 0.1025 (n=10000) | 0.0041 (n=10000) | 0.0012 (n=10000) |
| 0 | uniform | recessive | 0.089 (n=5000) | 0.08937 (n=9500) | 0.00027 (n=7500) | 0.00027 (n=7500) |
| 0 | uniform | additive | 0.1058 (n=5000) | 0.1038 (n=10000) | 0.0056 (n=10000) | 7e-04 (n=10000) |
| 0 | 0.1 | uniform | 0.129 (n=5000) | 0.131 (n=10000) | 0.0017 (n=10000) | 2e-04 (n=10000) |
| 0 | 0.1 | recessive | 0.09517 (n=6000) | 0.10129 (n=7000) | 0.00013 (n=7500) | 0 (n=6500) |
| 0 | 0.1 | additive | 0.1306 (n=5000) | 0.128 (n=10000) | 0.001 (n=10000) | 1e-04 (n=10000) |
| 0 | 0.5 | uniform | 0.1128 (n=5000) | 0.0997 (n=10000) | 0.0051 (n=10000) | 4e-04 (n=10000) |
| 0 | 0.5 | recessive | 0.095 (n=5000) | 0.0885 (n=10000) | 4e-04 (n=7500) | 0.00031 (n=6500) |
| 0 | 0.5 | additive | 0.1028 (n=5000) | 0.1037 (n=10000) | 0.0049 (n=10000) | 4e-04 (n=10000) |
| 0 | 1 | uniform | 0.0844 (n=5000) | 0.0875 (n=10000) | 0.0049 (n=10000) | 0.001 (n=10000) |
| 0 | 1 | recessive | 0.0792 (n=5000) | 0.0817 (n=10000) | 0.00042 (n=9500) | 0.00012 (n=8500) |
| 0 | 1 | additive | 0.0874 (n=5000) | 0.077 (n=10000) | 0.0092 (n=10000) | 1e-04 (n=10000) |
| 0.2 | uniform | uniform | 0.0946 (n=5000) | 0.0885 (n=10000) | 0.0033 (n=10000) | 2e-04 (n=10000) |
| 0.2 | uniform | recessive | 0.1398 (n=5000) | 0.1539 (n=10000) | 0.00462 (n=6500) | 0 (n=9000) |
| 0.2 | uniform | additive | 0.0952 (n=5000) | 0.0901 (n=10000) | 0.0054 (n=10000) | 2e-04 (n=10000) |
| 0.2 | 0.1 | uniform | 0.1178 (n=5000) | 0.1211 (n=10000) | 0.0015 (n=10000) | 3e-04 (n=10000) |
| 0.2 | 0.1 | recessive | 0.14436 (n=5500) | 0.138 (n=6000) | 0.01154 (n=6500) | 0 (n=7000) |
| 0.2 | 0.1 | additive | 0.1164 (n=5000) | 0.1258 (n=10000) | 0.0028 (n=10000) | 2e-04 (n=10000) |
| 0.2 | 0.5 | uniform | 0.0912 (n=5000) | 0.089 (n=10000) | 0.0048 (n=10000) | 5e-04 (n=10000) |
| 0.2 | 0.5 | recessive | 0.154 (n=5000) | 0.1427 (n=10000) | 0.00322 (n=9000) | 0 (n=8500) |

| $p_{\text{self}}$ | s | h | All benefits | Compensation | No compensation | No benefit |
| --- | --- | --- | --- | --- | --- | --- |
| 0.2 | 0.5 | additive | 0.0884 (n=5000) | 0.0826 (n=10000) | 0.0041 (n=10000) | 5e-04 (n=10000) |
| 0.2 | 1 | uniform | 0.0808 (n=5000) | 0.0672 (n=10000) | 0.0055 (n=10000) | 4e-04 (n=10000) |
| 0.2 | 1 | recessive | 0.1428 (n=5000) | 0.1378 (n=10000) | 0.00056 (n=9000) | 0.00084 (n=9500) |
| 0.2 | 1 | additive | 0.0746 (n=5000) | 0.0669 (n=10000) | 0.0062 (n=10000) | 6e-04 (n=10000) |
| 0.4 | uniform | uniform | 0.0762 (n=5000) | 0.0736 (n=10000) | 0.004 (n=10000) | 2e-04 (n=10000) |
| 0.4 | uniform | recessive | 0.1598 (n=5000) | 0.1534 (n=10000) | 0.0086 (n=10000) | 0.00021 (n=9500) |
| 0.4 | uniform | additive | 0.0824 (n=5000) | 0.0738 (n=10000) | 0.0048 (n=10000) | 5e-04 (n=10000) |
| 0.4 | 0.1 | uniform | 0.105 (n=5000) | 0.109 (n=10000) | 0.0017 (n=10000) | 1e-04 (n=10000) |
| 0.4 | 0.1 | recessive | 0.131 (n=5000) | 0.1275 (n=10000) | 0.00778 (n=9000) | 0 (n=8500) |
| 0.4 | 0.1 | additive | 0.1086 (n=5000) | 0.103 (n=10000) | 0.0028 (n=10000) | 5e-04 (n=10000) |
| 0.4 | 0.5 | uniform | 0.071 (n=5000) | 0.0681 (n=10000) | 0.0053 (n=10000) | 3e-04 (n=10000) |
| 0.4 | 0.5 | recessive | 0.1672 (n=5000) | 0.1617 (n=10000) | 0.0107 (n=10000) | 9e-04 (n=10000) |
| 0.4 | 0.5 | additive | 0.0714 (n=5000) | 0.0718 (n=10000) | 0.0083 (n=10000) | 5e-04 (n=10000) |
| 0.4 | 1 | uniform | 0.0542 (n=5000) | 0.0547 (n=10000) | 0.0066 (n=10000) | 0 (n=10000) |
| 0.4 | 1 | recessive | 0.1468 (n=5000) | 0.149 (n=10000) | 5e-04 (n=10000) | 4e-04 (n=10000) |
| 0.4 | 1 | additive | 0.0626 (n=5000) | 0.0532 (n=10000) | 0.0074 (n=10000) | 8e-04 (n=10000) |
| 0.6 | uniform | uniform | 0.0576 (n=5000) | 0.0578 (n=10000) | 0.0043 (n=10000) | 7e-04 (n=10000) |
| 0.6 | uniform | recessive | 0.0702 (n=5000) | 0.0672 (n=10000) | 0.0013 (n=10000) | 7e-04 (n=10000) |
| 0.6 | uniform | additive | 0.0618 (n=5000) | 0.058 (n=10000) | 0.0057 (n=10000) | 5e-04 (n=10000) |
| 0.6 | 0.1 | uniform | 0.0886 (n=5000) | 0.0917 (n=10000) | 0.0021 (n=10000) | 3e-04 (n=10000) |
| 0.6 | 0.1 | recessive | 0.0938 (n=5000) | 0.1021 (n=10000) | 2e-04 (n=10000) | 5e-04 (n=10000) |
| 0.6 | 0.1 | additive | 0.0906 (n=5000) | 0.091 (n=10000) | 0.0036 (n=10000) | 2e-04 (n=10000) |
| 0.6 | 0.5 | uniform | 0.0546 (n=5000) | 0.0512 (n=10000) | 0.006 (n=10000) | 7e-04 (n=10000) |
| 0.6 | 0.5 | recessive | 0.0634 (n=5000) | 0.0607 (n=10000) | 0.0031 (n=10000) | 6e-04 (n=10000) |
| 0.6 | 0.5 | additive | 0.0594 (n=5000) | 0.0557 (n=10000) | 0.006 (n=10000) | 2e-04 (n=10000) |
| 0.6 | 1 | uniform | 0.0488 (n=5000) | 0.0399 (n=10000) | 0.0035 (n=10000) | 8e-04 (n=10000) |
| 0.6 | 1 | recessive | 0.048 (n=5000) | 0.0417 (n=10000) | 4e-04 (n=10000) | 6e-04 (n=10000) |
| 0.6 | 1 | additive | 0.0516 (n=5000) | 0.0445 (n=10000) | 0.0053 (n=10000) | 4e-04 (n=10000) |
| 0.8 | uniform | uniform | 0.0374 (n=5000) | 0.0433 (n=10000) | 0.0035 (n=10000) | 4e-04 (n=10000) |
| 0.8 | uniform | recessive | 0.042 (n=5000) | 0.0455 (n=10000) | 0.0023 (n=10000) | 2e-04 (n=10000) |
| 0.8 | uniform | additive | 0.042 (n=5000) | 0.0416 (n=10000) | 0.0026 (n=10000) | 5e-04 (n=10000) |
| 0.8 | 0.1 | uniform | 0.0658 (n=5000) | 0.0665 (n=10000) | 0.0022 (n=10000) | 5e-04 (n=10000) |
| 0.8 | 0.1 | recessive | 0.0678 (n=5000) | 0.0747 (n=10000) | 7e-04 (n=10000) | 4e-04 (n=10000) |
| 0.8 | 0.1 | additive | 0.066 (n=5000) | 0.066 (n=10000) | 0.0018 (n=10000) | 4e-04 (n=10000) |
| 0.8 | 0.5 | uniform | 0.0386 (n=5000) | 0.0354 (n=10000) | 0.0042 (n=10000) | 0.0012 (n=10000) |
| 0.8 | 0.5 | recessive | 0.0366 (n=5000) | 0.0323 (n=10000) | 0.0023 (n=10000) | 3e-04 (n=10000) |
| 0.8 | 0.5 | additive | 0.0404 (n=5000) | 0.0375 (n=10000) | 0.0034 (n=10000) | 1e-04 (n=10000) |
| 0.8 | 1 | uniform | 0.0308 (n=5000) | 0.0301 (n=10000) | 0.0032 (n=10000) | 3e-04 (n=10000) |
| 0.8 | 1 | recessive | 0.033 (n=5000) | 0.0285 (n=10000) | 4e-04 (n=10000) | 5e-04 (n=10000) |
| 0.8 | 1 | additive | 0.0388 (n=5000) | 0.0322 (n=10000) | 0.0041 (n=10000) | 2e-04 (n=10000) |
| 1 | uniform | uniform | 0.0132 (n=5000) | 0.0138 (n=10000) | 0.0011 (n=10000) | 6e-04 (n=10000) |
| 1 | uniform | recessive | 0.0102 (n=5000) | 0.0072 (n=10000) | 8e-04 (n=10000) | 2e-04 (n=10000) |
| 1 | uniform | additive | 0.013 (n=5000) | 0.0127 (n=10000) | 0.0013 (n=10000) | 4e-04 (n=10000) |
| 1 | 0.1 | uniform | 0.0242 (n=5000) | 0.0223 (n=10000) | 4e-04 (n=10000) | 2e-04 (n=10000) |

| $p_{\text{self}}$ | $s$ | $h$ | All benefits | Compensation | No compensation | No benefit |
| --- | --- | --- | --- | --- | --- | --- |
| 1 | 0.1 | recessive | 0.0214 (n=5000) | 0.0235 (n=10000) | 9e-04 (n=10000) | 8e-04 (n=10000) |
| 1 | 0.1 | additive | 0.0196 (n=5000) | 0.0151 (n=10000) | 5e-04 (n=10000) | 3e-04 (n=10000) |
| 1 | 0.5 | uniform | 0.0148 (n=5000) | 0.0154 (n=10000) | 0.0021 (n=10000) | 6e-04 (n=10000) |
| 1 | 0.5 | recessive | 0.012 (n=5000) | 0.0084 (n=10000) | 0.0028 (n=10000) | 5e-04 (n=10000) |
| 1 | 0.5 | additive | 0.0168 (n=5000) | 0.0161 (n=10000) | 0.0023 (n=10000) | 2e-04 (n=10000) |
| 1 | 1 | uniform | 0.023 (n=5000) | 0.0228 (n=10000) | 0.0023 (n=10000) | 6e-04 (n=10000) |
| 1 | 1 | recessive | 0.0268 (n=5000) | 0.0267 (n=10000) | 8e-04 (n=10000) | 6e-04 (n=10000) |
| 1 | 1 | additive | 0.0246 (n=5000) | 0.025 (n=10000) | 0.0027 (n=10000) | 5e-04 (n=10000) |

**Figure S1:**
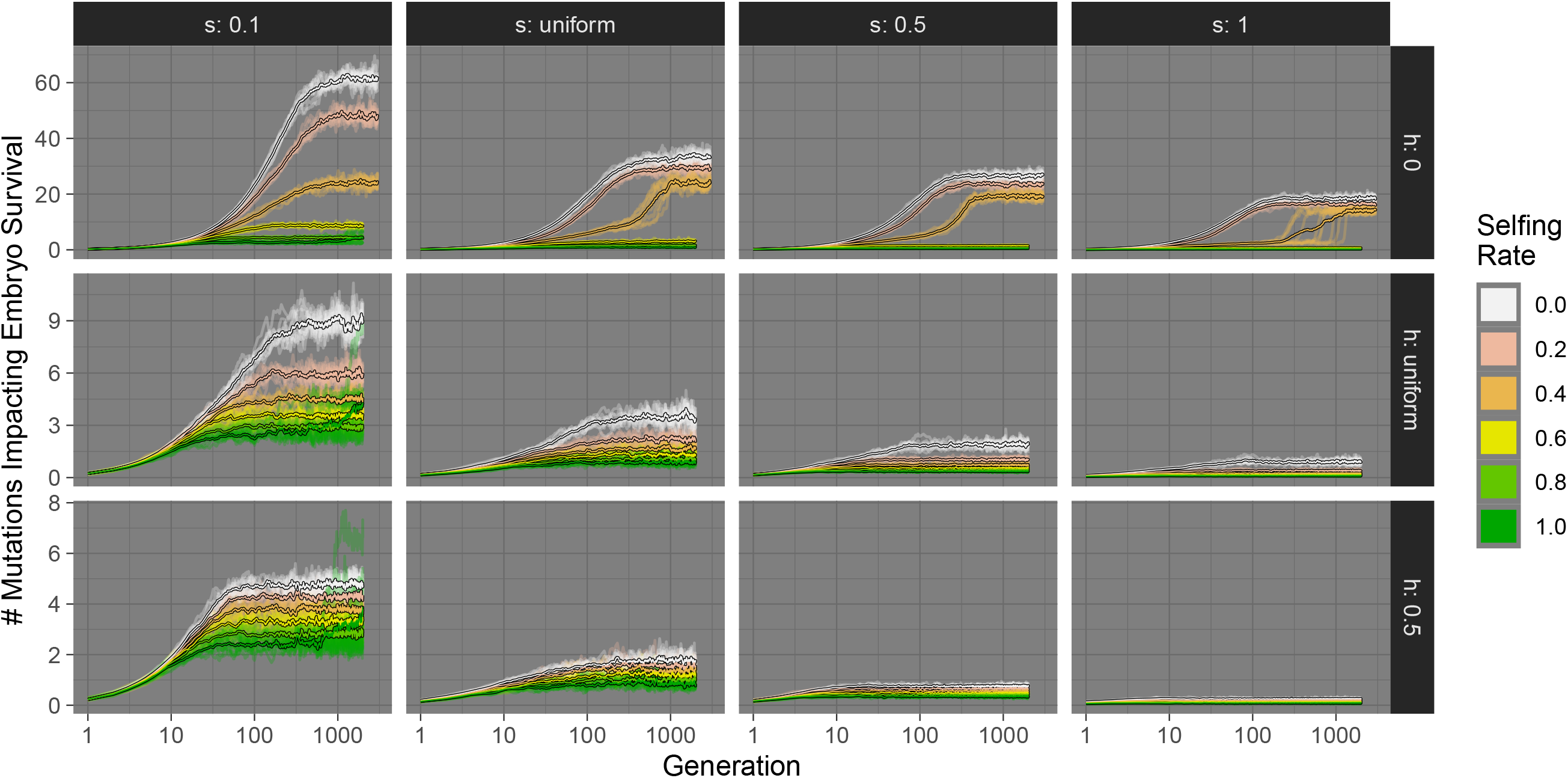
The accumulation of mutations: The number of deleterious mutations impacting embryo fitness over time in burn-in simulations, across selective (*s*) and dominance (*h*) coefficients, and selfing rates (on the x-axis).

**Figure S2:**
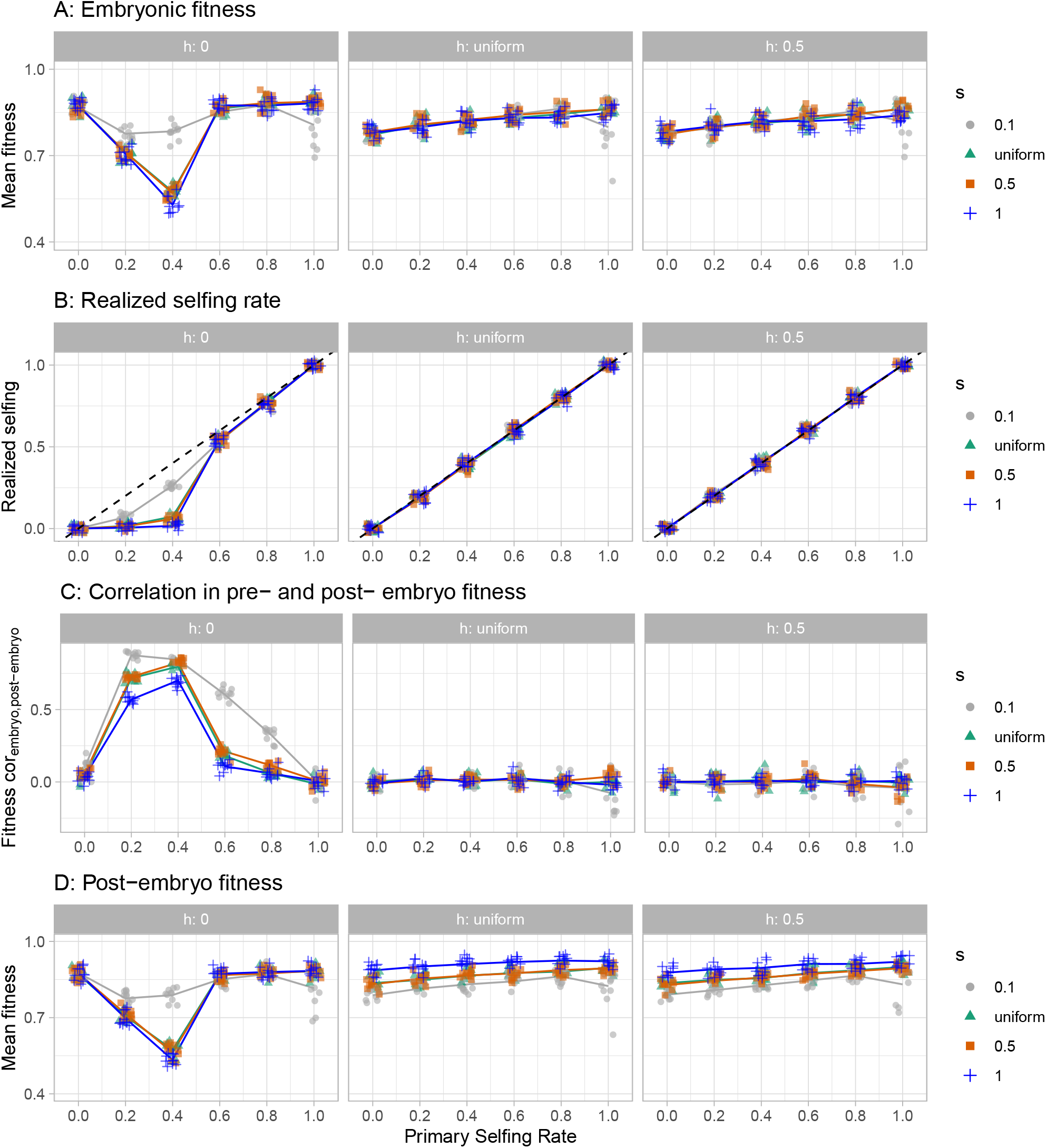
Summaries of our simulated populations at equilibrium: **(A)** The mean embryo fitness, **(B)** Realized selfing rate, **(C)** Correlation between embryo and postembryonic fitness, **(D)** Post-embryo fitness, across selective (*s*, colors and shapes) and dominance (*h*, facets) coefficients, and selfing rates (on the x-axis). The number of deleterious mutations impacting embryo fitness over time in burn-in simulations, across selective (*s*) and dominance (*h*) coefficients, and selfing rates (on the x-axis).

**Figure S3:**
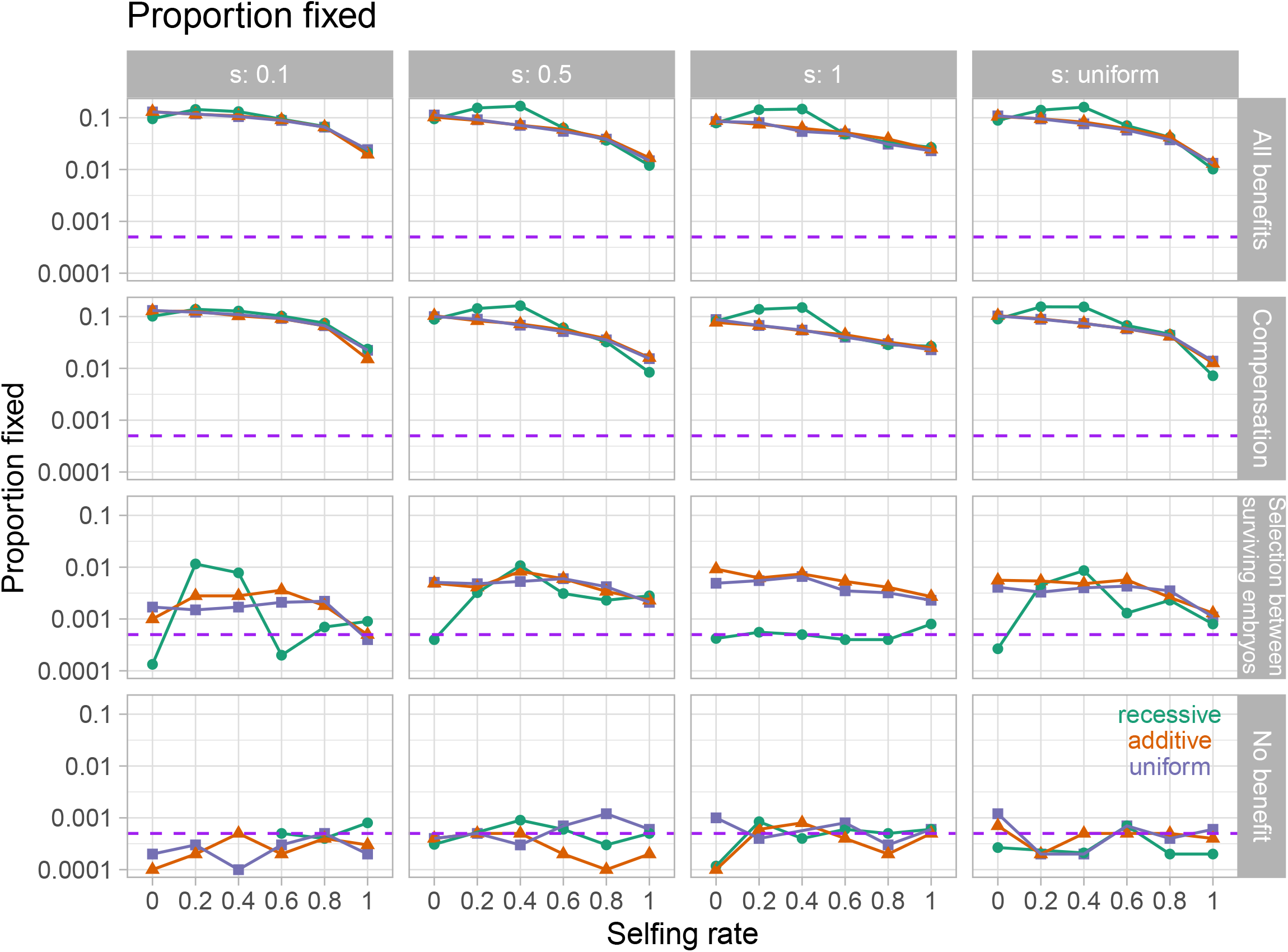
Proportion of introductions resulting in fixation as a function of selfing rate (x-axis), the benefit of polyembryony (rows), selection against new mutations (columns), and the dominance of new mutations (color). The purple line denotes neutral expectations. Note that fixation probabilities for additive mutations and those taking their value from a uniform distribution are very similar.

**Figure S4:**
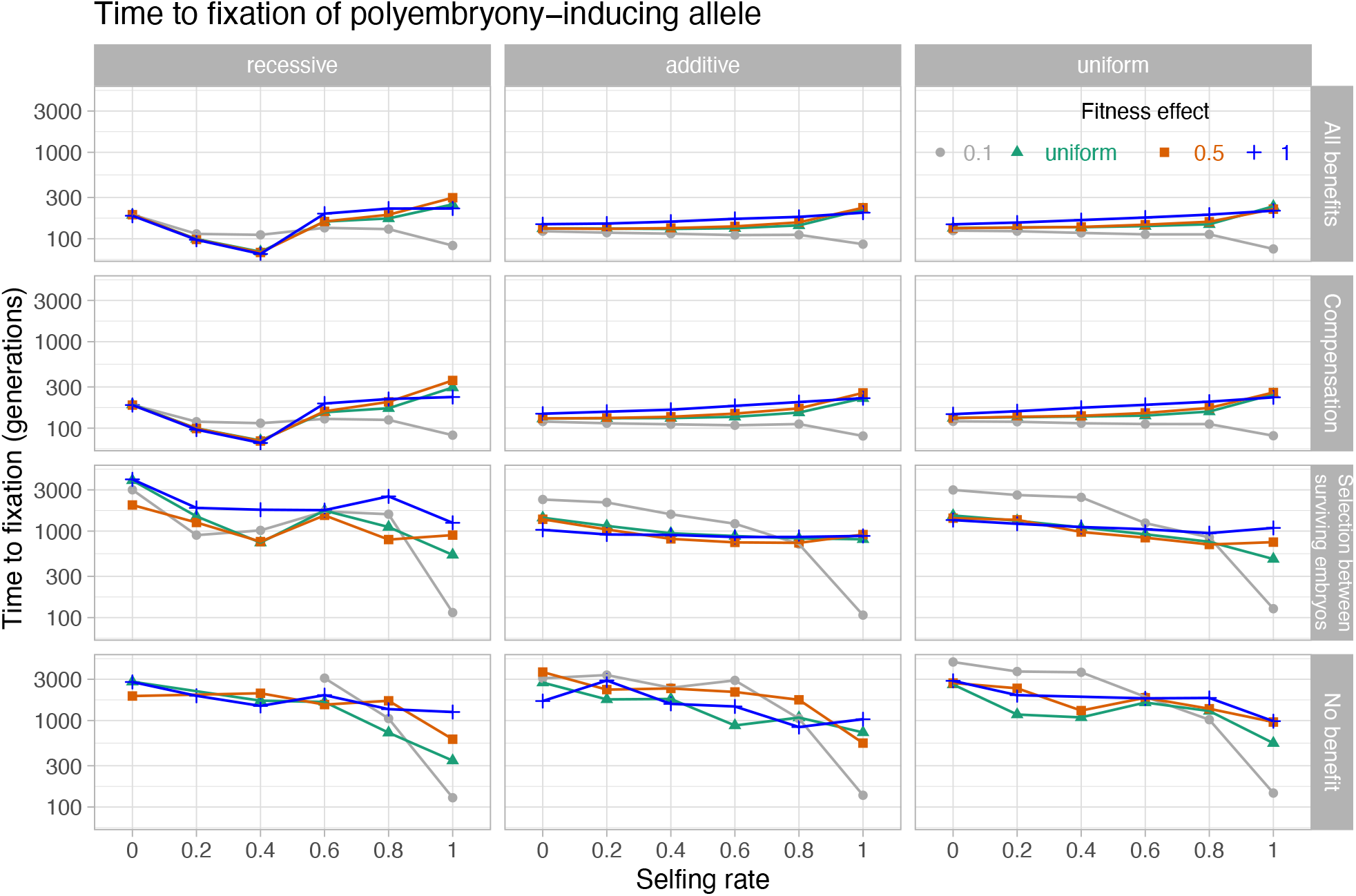
Mean time to fixation of the polyembryony allele across selective (s, colors and shapes) and dominance (h, faceted columns) coefficients, and selfing rates (on the x-axis), for each model (faceted rows).

**Figure S5:**
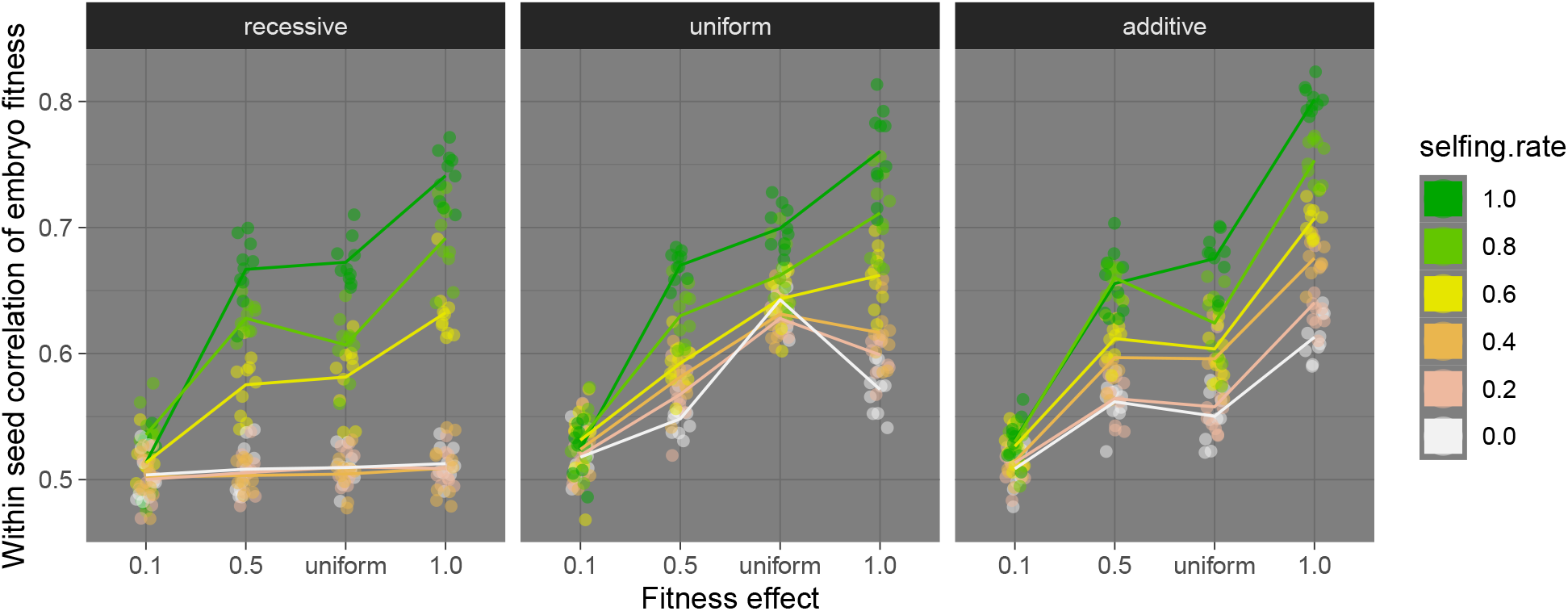
Correlation in fitness of the hypothetical two embryos in a seed, before polyembryony evolves as a function of the fitness effect of new mutations (on the x-axis), the selfing rate (color), and the dominance effect of new mutations (columns).

**Figure S6:**
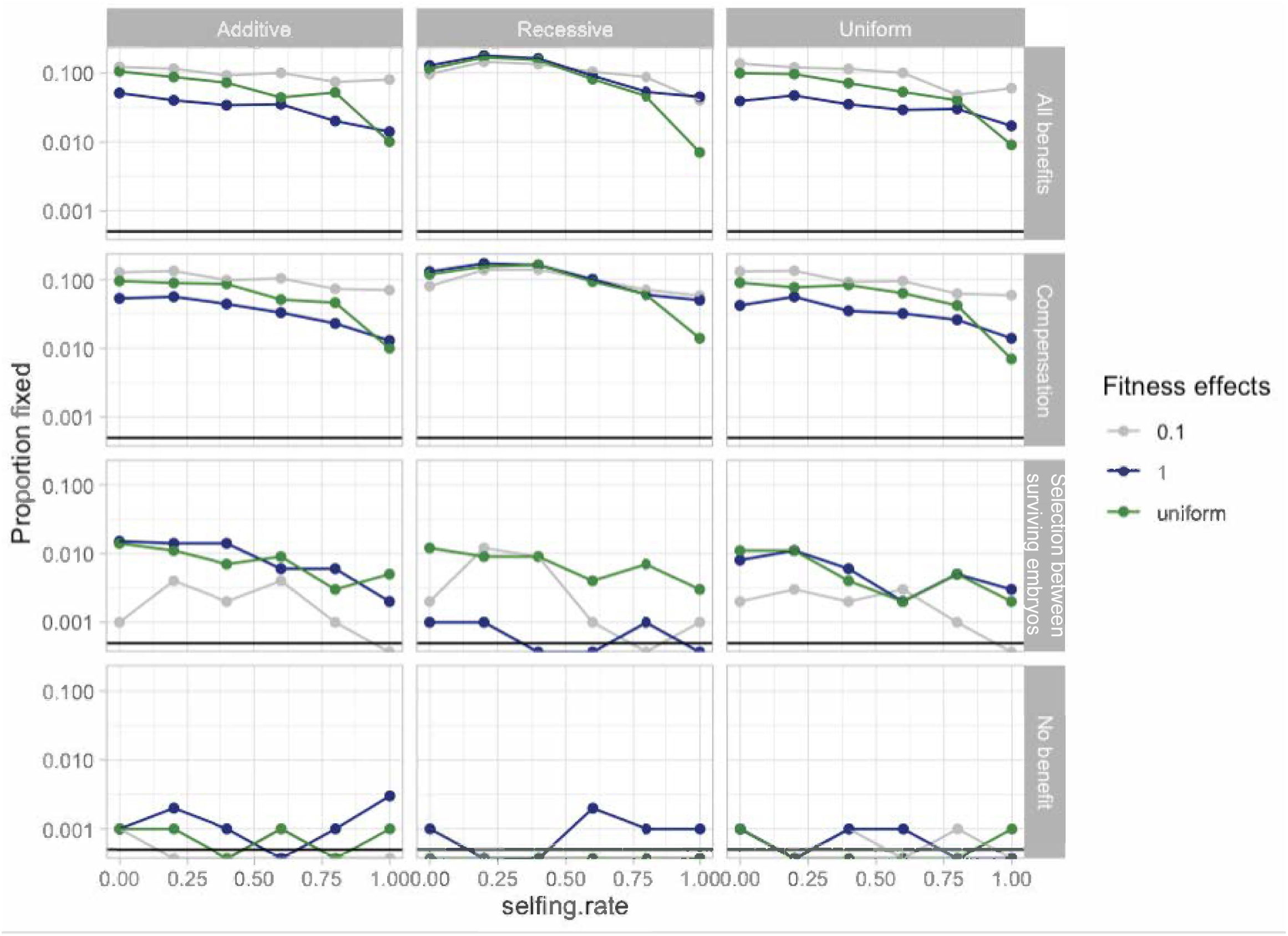
Proportion of introductions resulting in fixation with complete pleiotropy as a function of selfing rate (x-axis), the benefit of polyembryony (rows), selection against new mutations (columns), and the dominance of new mutations (color). The black line denotes neutral expectations. In this model every mutations impacts both embryonic and post-embryonic fitness in the same way.

## Notes

### Competing Interest Statement

The authors have declared no competing interest.

### Summary of Updates

We have edited the manuscript through for clarity. The most substantive changes revolve around clarity of what the model with selection between surviving embryos model -- highlighting that this "control" model is provided as a theoretical tool to isolate the two potential benefits of polyembryony, and not a model grounded in a particular biological process.

